# Molecular evolutionary dynamics of energy limited microorganisms

**DOI:** 10.1101/2021.02.08.430186

**Authors:** William R. Shoemaker, Evgeniya Polezhaeva, Kenzie B. Givens, Jay T. Lennon

## Abstract

Microorganisms have the unique ability to survive extended periods of time in environments with extremely low levels of exploitable energy. To determine the extent that energy limitation affects microbial evolution, we examined the molecular evolutionary dynamics of a phylogenetically diverse set of taxa over the course of 1,000-days. We found that periodic exposure to energy limitation affected the rate of molecular evolution, the accumulation of genetic diversity, and the rate of extinction. We then determined the degree that energy limitation affected the spectrum of mutations as well as the direction of evolution at the gene level. Our results suggest that the initial depletion of energy altered the direction and rate of molecular evolution within each taxon, though after the initial depletion the rate and direction did not substantially change. However, this consistent pattern became diminished when comparisons were performed across phylogenetically distant taxa, suggesting that while the dynamics of molecular evolution under energy limitation are highly generalizable across the microbial tree of life, the targets of adaptation are specific to a given taxon.

## Introduction

Organisms frequently encounter environments where growth and reproduction cannot be maintained. This is particularly true for microorganisms, where growth and reproduction is often limited by the flux of exploitable energy^1–3^. The occurrence of microorganisms in these environments is frequent enough that it has been argued that energy limitation constitutes a general constraint on microbial metabolism and growth^4–6^. Because evolution is fundamentally a birth-death process^7^, if a microbial population does not go extinct the extent that energy limitation constrains reproduction would in turn affect its subsequent evolution^8^. However, the majority of evolution experiments are performed in environments that permit rapid growth and those that focus on energy limitation are conducted with a single taxon^9–13^, limiting our ability to draw general conclusions about the evolutionary effects of energy limitation.

In order to understand how energy limitation alters microbial evolutionary dynamics it is necessary to first identify patterns that consistently occur across taxa. These patterns include the rate that *de novo* mutations accumulate, their nucleotide spectra, and how they change in frequency over time. Given that the majority of mutations are introduced during genome replication^14^, once cells can no longer reproduce the input of genetic variation should effectively cease until environmental conditions improve and reproduction can resume. However, this prediction can be violated if microorganisms in energy limited environments continue to reproduce at an extremely low rate^9,10^, a phenomenon known as “cryptic growth”^15^, where genetic diversity can accumulate and environmental effects on the nucleotide spectrum can occur^8,16–20^.

However, the evolutionary effect of energy limitation is not limited to the quantity and rate of change of genetic diversity. Rather, certain genes repeatedly acquire mutations in energy limited environments^10,12,21–26^. These repeated evolutionary outcomes (i.e., parallel evolution) consistently occur across phylogenetically distant taxa^27–32^, suggesting that adaptation continues as the environment becomes increasingly depleted of exploitable energy. Though such studies rarely manipulate the degree of energy limitation, contrast their observations to those from populations in energy replete environments, or perform comparisons across multiple taxa. These shortcomings prevent us from determining whether the targets of molecular evolution change as the environment transitions from an energy replete to depleted state (i.e., divergent evolution) as well as whether any such change in the direction of evolution can be generalized across the microbial tree of life.

In this study we examined the molecular evolutionary dynamics of energy limited microbial populations from six bacterial taxa that were propagated for ∼1,000 days, which were chosen to maximize phylogenetic diversity subject to the condition that they all grew in the same environmental conditions (*Bacillus, Caulobacter, Deinococcus, Janthinobacterium, Pedobacter*, and *Pseudomonas*). We manipulated energy limitation by extending the time between transfer events, corresponding to transfer times of 1, 10, and 100 days, which allowed us to examine evolutionary outcomes across a range of energy limitation regimes as well as taxa. We examined how genetic diversity, the spectrum of mutations, and the distribution of extinction events were affected by energy limitation. We then examined the degree that energy limitation affected the direction of molecular evolution across all taxa. Finally, we determined whether phylogenetically diverse taxa converged on similar molecular evolutionary outcomes within and across energy limitation regimes.

## Results

Molecular evolutionary dynamics were consistently affected by transfer time, our proxy for energy limitation, across phylogenetically diverse taxa. In energy depleted environments, mutations accumulated at a faster rate than expected based on the timescale of the experiment across all taxa, though the molecular evolutionary dynamics of 10-day and 100-day regimes were more similar than either were to the 1-day regime. We found that the spectrum of mutations was consistently affected by energy limitation across all taxa, though the effect on the proportion of nonsynonymous and synonymous polymorphisms (*pN/pS*) was less consistent, suggesting that the strength of negative selection did not change as energy limitation increased. By comparing the genes that acquired excess nonsynonymous mutations, we found a consistent pattern of divergent evolution among all energy limitation regimes and taxa, where, again, the 10 and 100-day transfer regimes had the highest degree of similarity. However, this pattern of divergent evolution was absent when comparisons were made at a coarser phylogenetic scale, suggesting that it is unlikely that the targets of molecular evolution under energy limitation within a taxon can be generalized across the tree of life. The recurrent patterns we observed within a given taxon lead us to conclude that the rate and direction of molecular evolution consistently shifted across all taxa after energy depletion occurred, but remained relatively constant thereafter.

### Dynamics and fates of mutations under energy limitation

Microbial populations continue to acquire genetic diversity under energy limited conditions^10^, though their quantitative dynamics are rarely examined. To address this, we constructed a molecular fossil record for ∼100 populations that evolved for ∼1,000 days (Figs. S1-6). By examining the accumulation of mutations by time *t* as the sum of derived allele frequencies (*M*(*t*)), we found that the accumulation of mutations was fairly consistent across energy limitation regimes, where *M*(*t*) typically saturated by day 500 (Figs. S7-12,a-c). We found that the change in *M*(*t*) between timepoints (Δ*M*(*t*)) was also consistent across energy limitation regimes and taxa (Figs. S7-12,d-f), suggesting that these general molecular evolutionary patterns were not affected by energy limitation.

By examining the relationship between *M*(*t*) and the degree of energy limitation, we were able to determine the degree that different taxa continued to acquire mutations under energy limitation. If cell division completely stopped once the environment was exhausted of energy and the majority of *de novo* mutations were acquired due to genome replication errors, then the number of mutations *M*(*t*) should decay as the time between transfers increased. All taxa deviated from this prediction, where the slope was consistently greater than the null expectation (*β*_1_ *>* −1; Fig. 1a-f). This deviation was extreme in certain cases, as the slopes of *Pseudomonas* and *Pedobacter* were virtually zero, indicating that the amount of accumulated genetic diversity remained highly similar despite the wide range of energy limitation regimes. Conversely, the slope of *Bacillus* was closest to the null expectation.

**Figure 1.**
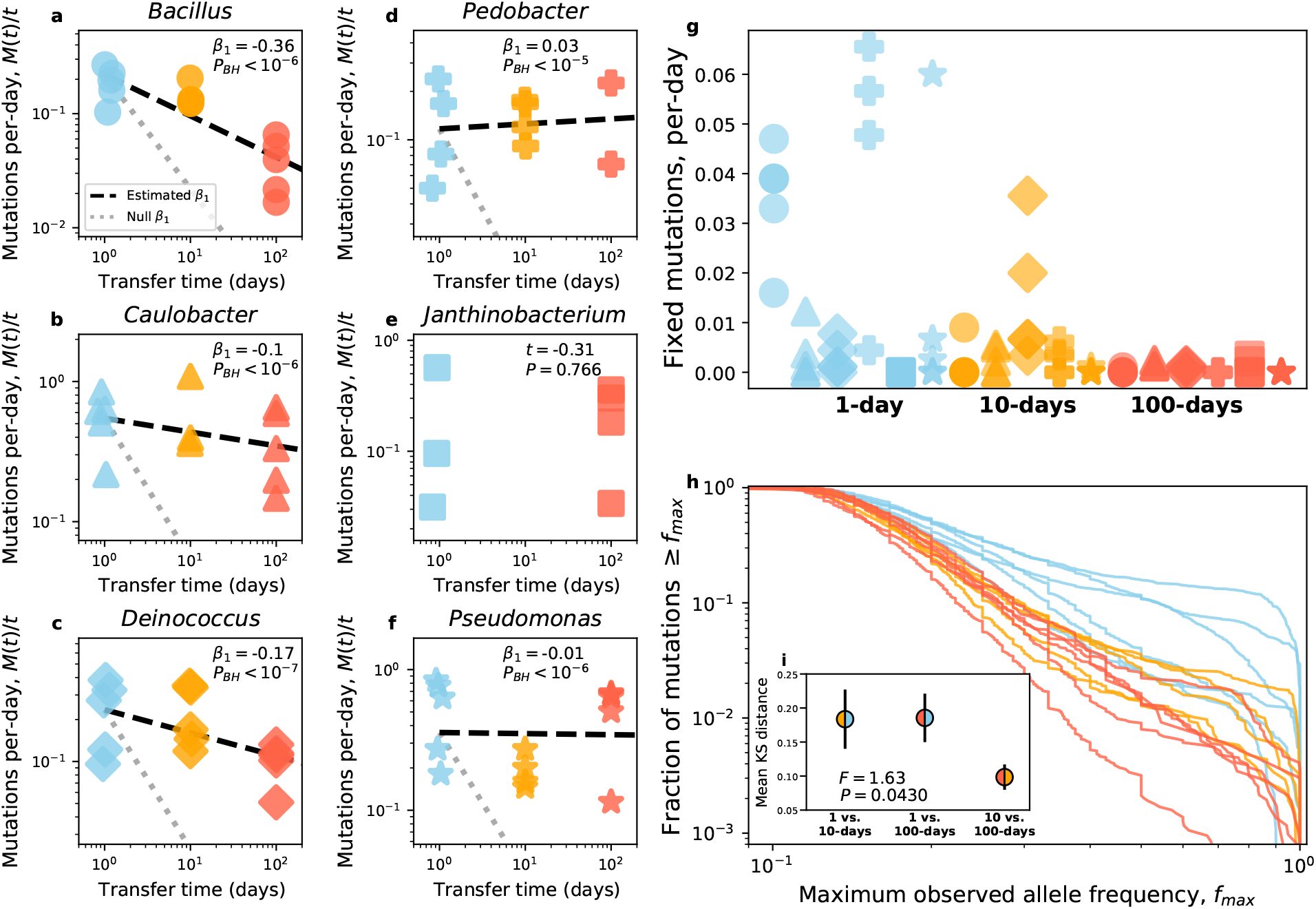
**a-f)** The relationship between mutations and transfer time varied across taxa and deviated from the null expectation (grey dotted line). However, the slope of *Bacillus* was the closest to the null expectation, consistent with the interpretation that dormancy acted as an evolutionary buffer. A *t*-test was performed on *Janthinobacterium* instead of a regression due to the high extinction rate of populations in the 10-day energy limitation regime. **g)** The total number of fixation events was fairly low and highly variable regardless of treatment. **h)** As an alternative analysis, we examined the empirical cumulative distribution of the maximum frequency that mutations reached *f*_*max*_ for each taxon-treatment combination. To examine which energy limitation regimes were similar, we calculated the mean Kolmogorov-Smirnov distance among all taxa for a given pair of energy limitation regimes (inset figure in **h**, black bars represent the standard error of the mean). Samples from a Gaussian have been added to the x-axes of **a-f** for visibility.

Surprisingly, very few mutations appeared to fix over the course of 1,000 days, even in populations that were transferred daily (Figs. S1-6). Using a previously published Hidden Markov Model-based approach^33^, we confirmed this observation by assigning each mutation a final state (i.e., fixed, extinct, or polymorphic; Figs. 1g, S7-12). The comparative absence of fixations made it difficult to compare substitution rates between treatments and taxa due to the excess number of zeros. However some general conclusions could still be made. Beyond the low number of fixation events, only *Pedobacter* and *Bacillus* experienced a sharp decline in fixation events as the time between transfers increased (Fig. 1g). While there is no obvious explanation for the disparity in the former, the sharp decline in *Bacillus* was likely driven, again, by its ability to form protective endospores.

Despite the lack of fixations, it was clear that few mutations reached frequencies ≳0.4 in 10 and 100-day transfer regimes, even if the change in *M*(*t*) remained constant (Figs. S7-12). Leveraging this observation, we examined the distribution of the maximum observed frequency of all mutations (*f*_*max*_) to infer whether the trajectories of mutations varied among energy limitation regimes. We found that the distribution of *f*_*max*_ in the 1-day transfer regime was clearly distinct from the 10 and 100-day regimes, while the latter two regimes largely overlap (Fig. 1h). Direct comparison of *f*_*max*_ cumulative distributions confirmed this, as the mean Kolmogorov–Smirnov distance between all combinations of energy limitation regimes was significant (permutational ANOVA: *F* = 1.63, *P* = 0.0430). The mean distance remained high for 1 versus 10-days and 1 versus 100-days comparisons, but notably dropped for 10 versus 100-days comparisons (Fig. 1h). This drop in *f*_*max*_ suggests that the trajectory of *de novo* mutations was altered after the initial depletion of exploitable energy, where nominally beneficial mutations could not continue to increase in frequency.

We note the absence of any segregation of intermediate-frequency mutations (i.e., clade formation) across all taxa and transfer regimes. This observation seemingly contrasts with the repeated emergence of quasi-stable clades in an experiment with similar energy limitation regimes conducted over a similar evolutionary timescale with *E. coli*^34,35^ as well as the proposal that the emergence of clades is a general feature of microbial evolution^36^. Though the lack of structure in our experiment was likely caused by differences in experimental design, as our experiment was performed in well-mixed flasks rather than test tubes with a lack of headspace^34,35^, meaning that there was no spatial structure that could promote the emergence of population genetic structure. Though experimental design does not entirely account for the lack of structure, as other evolution experiments using well-mixed flasks have documented the emergence of co-existing clades, the Long-term Evolution Experiment (LTEE) with *E. coli* being the most notable example^33^. However, the average timescale for the emergence of new clades in the LTEE was ∼16,000 generations, far longer than our longest timescale of ∼3,000 generations. This observation suggests that while energy limitation alone is insufficient to promote the emergence of multiple clades on short evolutionary timescales, we cannot rule out the possibility of clades eventually emerging if the experiment had continued.

### The molecular spectra of *de novo* mutations

We determined the degree that energy limitation affected the spectrum of mutations by examining the transition rates of all mutations at four-fold redundant sites. By performing dimensionality reduction on a population-by-spectra matrix (i.e., PCA), we found that replicate populations largely grouped by their energy limitation regime (Fig. S13). A permutational multivariate analysis of variance (PERMANOVA) test using a permutation structure that controlled for taxon identity confirmed this observation (*F* = 5.77, *P <* 10^−4^). By examining the factor loadings we found the nucleotide spectra that are correlated with the first and second principal components (Table S1). The transversion A: T → C: G was highly correlated with the first principle component 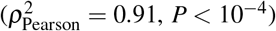 while the transition A: T → G: C was correlated with the second 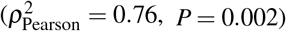, suggesting that these mutational spectra were affected by energy limitation.

Expanding our scope from synonymous mutations, we examined the spectrum of nonsynonymous and synonymous mutations to determine the degree that positive and negative selection change as energy limitation increases. Given the lack of fixation events within populations in 10 and 100-day transfer regimes, our comparative analyses were limited to polymorphic mutations. However, we were able to leverage the large number of mutations we observed to compare the ratio of nonsynonymous to synonymous polymorphisms (*pN/pS*) between transfer regimes, which we examined as a function of *f*_*max*_ rather than as a single snapshot so that we could control for the qualitatively different mutation trajectories found in each energy limitation regime. In general, we found that *pN/pS* remained considerably less than 1 as *f*_*max*_ increased, 1-day *Janthinobacterium* being the sole exception and only for *f*_*max*_ *>* 0.4 (Fig. 2e). This result is consistent with the hypothesis that negative selection predominantly occurs across microbial genomes^37,38^. To determine whether the relationship between *f*_*max*_ and *pN/pS* varied by energy limitation regime, we calculated the mean absolute deviation (MAD) for each taxon across all pairs of energy limitation regimes. We found that the mean absolute deviation in *pN/pS* was greater between the 1-day transfer regime and the 10 and 100-day transfer regimes 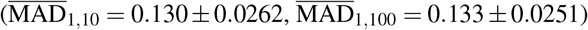 than for the latter two regimes 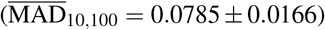. This pattern is consistent with the observation that the molecular evolutionary dynamics were affected by the initial depletion of energy and remained relatively constant as the degree of energy limitation increased. However, through a permutational ANOVA we found that the difference in 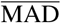 over our three treatment comparisons was not significant (*F* = 1.12, *P* = 0.446), suggesting that any change in the strength of negative selection as energy limitation increased was weak relative to the overall strength of negative selection operating across all regimes.

**Figure 2.**
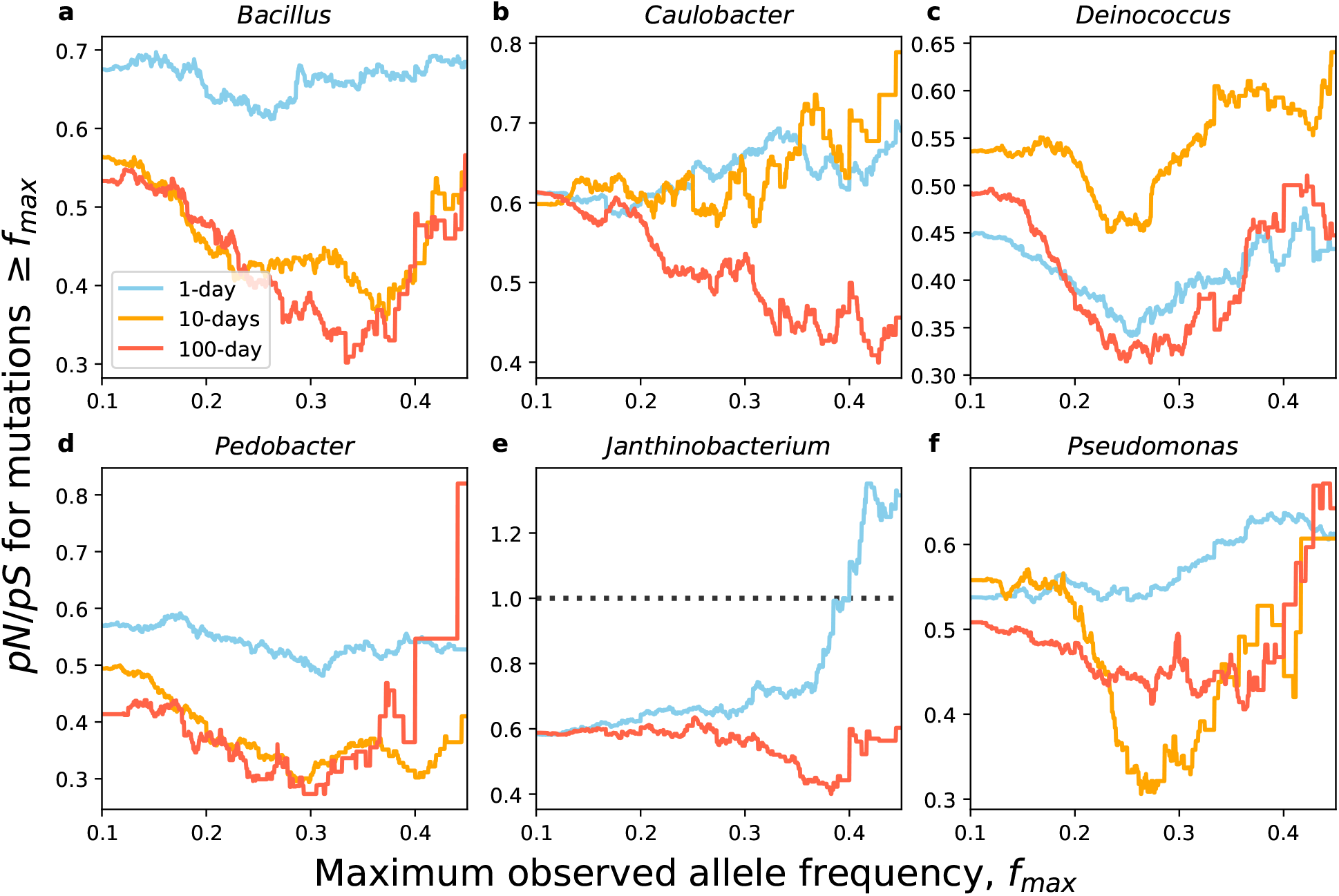
**a-f)** We examined the ratio of the number of nonsynonymous to synonymous polymorphic mutations contingent on the maximum frequency that a mutation reached (*f*_*max*_). Generally, values of *pN/pS* remain less than one across all transfer-regimes and taxa, suggesting the negative selection was highly prevalent. *Janthinobacterium* was the lone exception to this pattern, and only for high values of *f*_*max*_ within the 1-day treatment. The 10-day and 100-day *pN/pS* values tend to track one another, though *Caulobacter* and *Deinococcus* were notable exceptions. The black dashed line for **e** represents *pN/pS* = 1, the expectation under neutrality.

The majority of populations survived the experiment, though many *Caulobacter, Janthinobacterium*, and *Pedobacter* replicate populations repeatedly went extinct. Because all replicate populations were initially genetically identical, the dispersion of extinction events within a treatment contained information about whether the rate of extinction was contingent on a population’s evolutionary history. If the per-generation rate of extinction was sufficiently rare and an independent process, we would expect that the distribution of extinction events would follow a Poisson distribution. However, if mutations acquired by a given population over the course of the experiment affected the probability of extinction (i.e., historical contingency), the distribution should be overdispersed relative to a Poisson. By comparing the coefficient of variation calculated from empirical data (CV) to a null distribution generated from Poisson sampling, we found that extinctions in the 10-day transfer regime was overdispersed for *Pedobacter* and *Janthinobacterium* (Table 1). This result suggests that mutations acquired over the course of the experiment contributed to repeated extinction events in at least two taxa. It is worth noting that these two taxon-treatment combinations had the highest mean number of extinction events 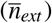, suggesting that historically contingent extinctions regularly occurred under energy limitation across phylogenetically distant taxa, but that an insufficient number of extinction events occurred in our experiment to detect this signal in all taxon-treatment combinations.

**Table 1.** The distribution of extinction events among replicate populations suggest historical contingency due to the acquisition of *de novo* mutations. By comparing the coefficient of variation (CV) of the number of extinction events to a null distribution calculated from simulations from a Poisson distribution generated using the mean number of extinction events 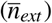, we determined whether historical contingency occurred within a given taxon-treatment combination. All taxon-treatment combinations where at least three replicate populations went extinct were examined.

| Taxon | Transfer time (d.) | $\bar{n}_{ext}$ | CV | $P_{BH}$ |
| --- | --- | --- | --- | --- |
| *Caulobacter* | 10 | 1.2 | 0.97 | 0.409 |
|  | 100 | 0.6 | 0.82 | 0.837 |
| *Pedobacter* | 10 | 5.4 | 0.74 | 0.03 |
|  | 100 | 3.2 | 0.31 | 0.837 |
| *Janthinobacterium* | 10 | 7.6 | 0.75 | 0.005 |
|  | 100 | 1.0 | 1.1 | 0.409 |

### Parallel, divergent, and convergent evolution within each taxon

To identify potential targets of adaptation, we examined the set of genes that acquired more nonsynonymous mutations than expected based on gene size within each energy limitation regime for each taxon (i.e., genes that contributed to parallel evolution). We first calculated the number of nonsynonymous mutations pooled across replicate populations at each gene after adjusting for gene size (i.e., multiplicity^33^) and tested whether nonsynonymous mutations were non-randomly distributed across genes (Supplementary Information). We easily rejected this null hypothesis for all taxon-treatment combinations (*P <* 10^−4^; Figs. S15-20, a). We note the presence of a small cluster of genes where the mean maximum observed allele frequency was fairly high 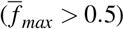, but multiplicity was low (Figs. S14-19, b). The formation of this cluster was likely driven by the fact that few mutations reached appreciable frequencies, which would decrease multiplicity given that it is weighted by total number of mutations in the population (Supplementary Information). However, few genes acquired more than one high frequency mutation, meaning that we were unable to identify the set of significantly enriched genes conditioned on a minimum *f*_*max*_. Instead, we determined whether parallelism at the gene level was dependent on allele frequency by selecting for mutations with a minimum value of *f*_*max*_ and calculating the likelihood difference of the enrichment of nonsynonymous mutations at certain genes across the genome versus no such enrichment (Δ*ℓ*). We found that Δ*ℓ* typically increased with *f*_*max*_ across energy limitation regimes for the majority of taxa (Fig. S14), which is consistent with the hypothesis that higher frequency mutations were primarily driven by positive selection across energy limitation regimes.

Using the set of genes that contributed to parallel evolution within each energy limitation regime for each taxon, for a given pair of energy limitation regimes we asked whether the evolved features of these genes were more similar or different than expected by chance. Operating under this broad definition of convergent and divergent evolution at the molecular level, we first examined whether the same genes were consistently enriched. Across taxa, the vast majority of genes that were enriched within a given treatment were enriched across all treatments (Figs. S21-25), which would suggest that convergent rather than divergent evolution occurred. Simulations modeled on the probability distribution describing the null expectation (Eq.1) confirmed this observation, as the proportion of genes that were enriched within both energy limitation regimes for a given pair was consistently greater than expected by chance for all pairwise combinations of energy limitation regimes across all taxa (Fig. 3a; Table S2). However, the degree of convergent evolution across pairs of energy limitation regimes was not equal, as the 10 versus 100-day comparison had a stronger degree of convergence than all other comparisons (Fig. 3a). This result builds on our above observations, providing evidence that the largest shift in the direction of evolution occurred shortly after the environment was depleted of exploitable energy.

**Figure 3.**
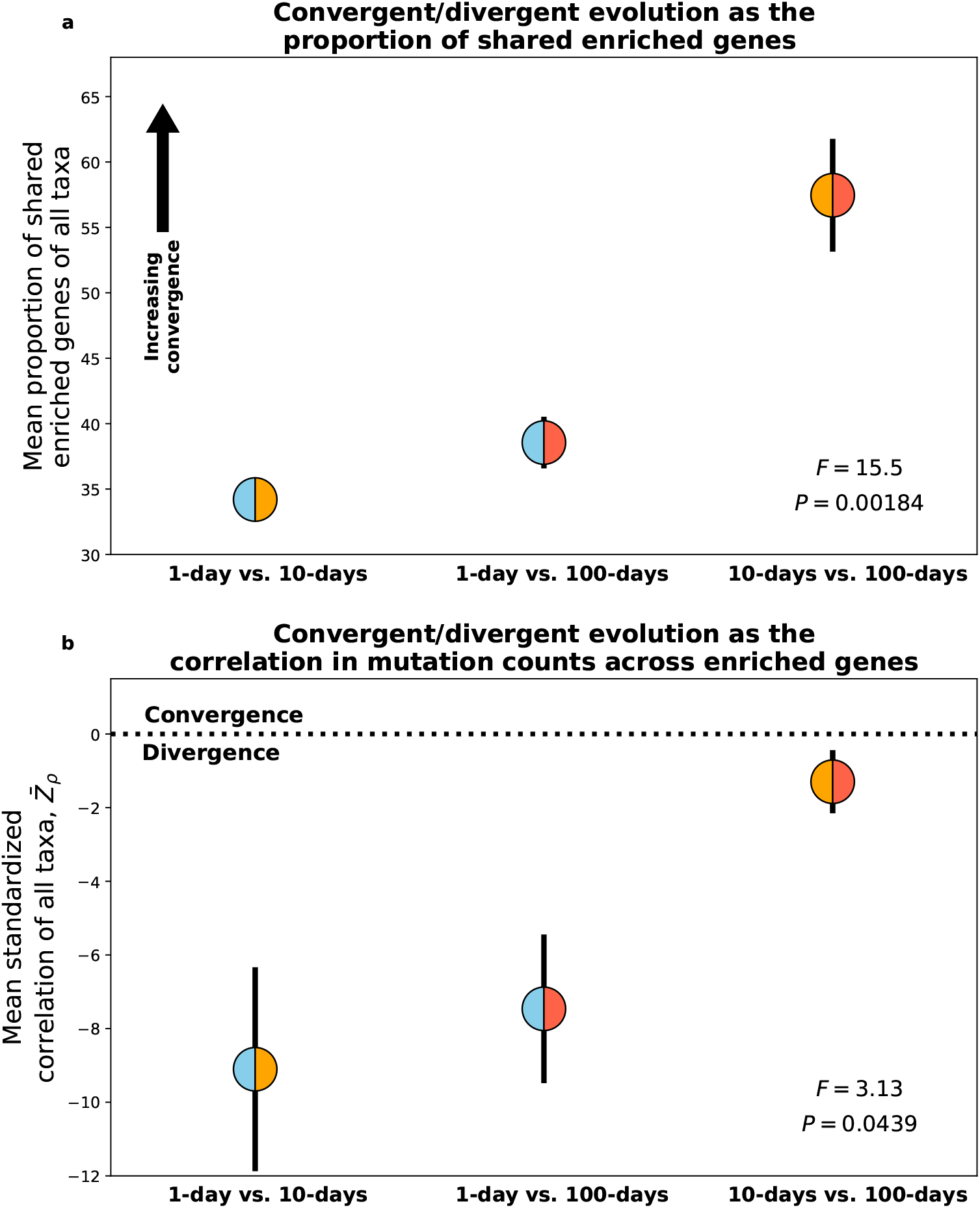
**a)** By defining convergent/divergent evolution as the degree that the proportion of genes that are enriched across treatments is greater or less than expected by chance, we find evidence of convergent evolution across the genome for all pairwise treatment combinations across taxa. A permutational ANOVA indicates that the degree of convergent evolution varied across treatments, the 10 and 100-day energy limitation regimes being the most similar. **b)** By examining the number of mutations within each gene, we can determine whether convergent or divergent evolution occurred within genes that were enriched for mutations by calculating the correlation coefficient between each pair of energy limitation regimes and standardizing it to a null distribution for each taxon. Here, standardized correlations less than zero correspond to divergent evolution while the opposite suggests convergence. In general, divergent evolution consistently occurred between all pairs of energy limitation regimes. While 10 and 100-day transfer regimes are more similar to each other than either are to the 1-day regime, the difference is borderline significant using a permutational ANOVA controlling for taxon identity. Black bars in both plots represent standard errors and each color scheme represents a given pair of energy limitation regimes.

While our definition of convergent/divergent evolution as the proportion of genes that were enriched across energy limitation regimes provided insight, it was likely too coarse an approach as it only considered gene identity. Assuming that each energy limitation manipulation acted as a slight perturbation on the rate of molecular evolution at each gene, gene identity alone would likely be insufficient to detect divergent evolution and would generally suggest convergent evolution of varying degrees. A direct comparison of mutation counts supports this claim, as the set of genes that were enriched across multiple energy limitation regimes consistently differed in the number of mutations acquired (Figs. S15-20, d-f). To quantify this potential divergence, we defined convergent and divergent evolution as the degree that the correlation of mutation counts between energy limitation regimes (*ρ*) was greater or less than expected by chance. To determine whether a given value of *ρ* constituted convergent or divergent evolution, we generated a null distribution by randomizing combinations of mutation counts while controlling for the total number of mutations acquired within each treatment and gene. By examining the standardized *ρ* (*Z*_*ρ*_) of each taxon, we found that divergent evolution consistently occurred across all combinations of energy limitation regimes (Fig. 3b). Similar to our analysis on gene identity, we found that the mean *Z*_*ρ*_ across taxa was closest to the null expectation for comparisons between 10 and 100-day transfers (Fig. 3g). The difference in the degree of divergent evolution was statistically significant (*F* = 3.13, *P* = 0.0439) and what we observed was consistent with the reoccurring pattern of the evolutionary dynamics in the 10 and 100-day energy limitation regimes having greater similarity to one another than either one did to the 1-day regime.

### Divergent and convergent evolution across taxa

While divergent and convergent evolution could readily be examined among populations within the same taxon, a similar analysis could not be performed among all phylogenetically diverse taxa due to the lack of shared gene content. Instead, it was necessary to map genes that were enriched for mutations to higher levels of biological organization that were present in all taxa. Therefore, we mapped genes to their functional modules using the Metabolic And Physiological potentiaL Evaluator (MAPLE)^39^, where we found several modules that were consistently hit across three or more taxa. To test whether this observation was significant, we simulated random draws of genes that were then mapped to their respective modules, from which we generated null rank curves for the number of modules that were hit across all possible combinations of taxa. We found that the observed decay curve deviated from our null expectation (Fig. 4a), but only up until four taxa for 1-day and 100-day transfers. This pattern suggests that convergent evolution did occur among taxa within the limits of energy repletion (1-day) and depletion (100-days), though the ability to detect this convergence dissipated as additional taxa were included. This reduced rate of decay was driven, at least in-part, by shared evolutionary history, as the similarity in enriched MAPLE modules decays with phylogenetic distance for 1 and 100-day transfer regimes (Fig. 4d). We note that while this trend was only significant for 1-day transfers, the slope for the 10-day treatment was virtually zero and the lack of a significant slope in the 100-day treatment can be explained by the low number of nonsynonymous mutations acquired by populations within that treatment, limiting our ability to identify enriched MAPLE modules and inflating the number of observations with zero overlap.

**Figure 4.**
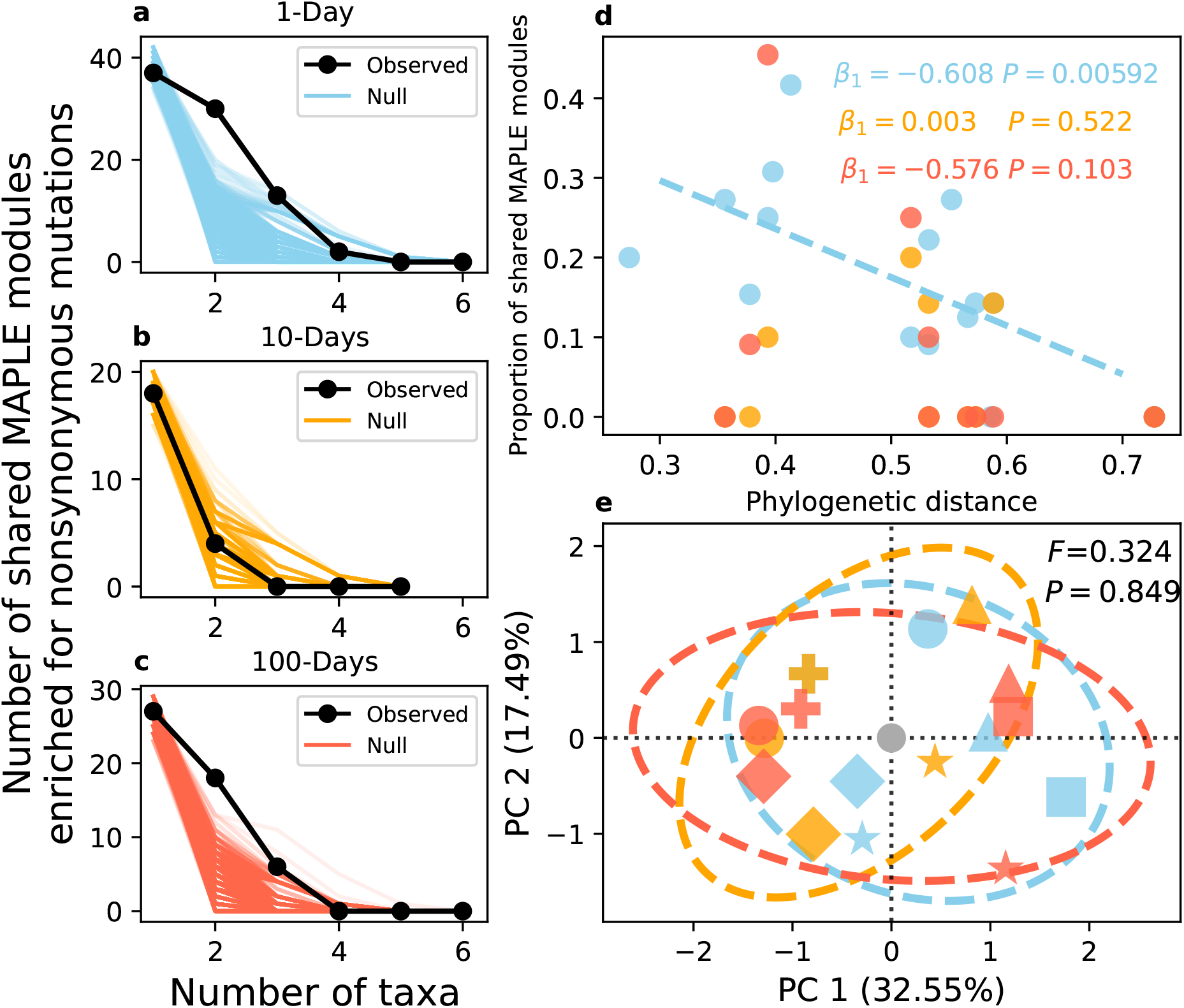
**a-c)** By comparing the observed decay of the number of MAPLE modules as the number of taxa increases to the null expectation, we can see that observed curve tended to decay at a slower rate than the null for 1 and 100-day transfer regimes, suggesting that convergent evolution occurred in those treatments. **d)** By examining the relationship between the degree of overlap in the set of enriched MAPLE modules (i.e., the Jaccard index) and the phylogenetic distance of a given pair of taxa, we then examined whether the degree of molecular convergent evolution was correlated with the amount of shared evolutionary history. The distance-decay relationship was significant for the 1-day transfer regime and was quite strong, though ultimately not significant for the 100-day regime, while the slope was virtually zero for the 10-regimes. This result suggests that the convergent evolution illustrated in **a** and **c** was driven by shared evolutionary history, where convergent evolution was more likely among phylogenetically similar taxa. **e)** PCA was performed to determine whether different MAPLE modules were enriched across different treatment groups (i.e., divergent evolution). Generally data points tended to cluster together by taxon identity rather than treatment. This result suggests that the signal of divergent evolution between energy limitation regimes that we observed in Fig. 3 did not persist at a coarse-grained functional level, a conclusion that is supported by the fact that effect of treatment was not significant via PERMANOVA.

While there were consistent patterns of convergent evolution across phylogenetically diverse taxa within the 1-day and 100-day energy limitation regimes, there remained the question of whether each regime acquired nonsynonymous mutations in a distinct set of modules. In other words, it was necessary to determine whether energy limitation continued to affect the direction of molecular evolution when all taxa are examined together instead of on an individual basis. By constructing a module-by-taxon/treatment matrix and performing PCA, we found that taxon-treatment combinations tended to cluster together by taxon identity rather than their energy limitation regime (Fig. 4e). A PERMANOVA test supports this observation (Fig. 4e). While there was no evidence of genome-wide divergent evolution between energy limitation regimes when comparisons were made at a higher phylogenetic level, we identified individual modules that may be of interest to future research efforts (Table 2). Overall there was a high degree of overlap among energy limitation regimes, where Coenzyme A biosynthesis was the only module enriched in at least two taxa within a single transfer regime. Iron complex transport systems also appeared to be slightly more enriched in 100-day transfer regimes, a micronutrient that has been shown to be essential for the survival of microorganisms in energy limited conditions^30,40^.

**Table 2.** The number of taxa with genes within a MAPLE module that were enriched for nonsynonymous mutations. Only modules that were enriched in at least two taxa within a given treatment regime are included in this table. There is a large amount of overlap between transfer-regimes, suggesting that highly conserved genes contributed towards adaptation in similar ways in energy deplete and replete environments.

| Module ID | Module type | Description | Number of taxa enriched/ Total number of taxa |  |  |
| --- | --- | --- | --- | --- | --- |
|  |  |  | 1-day | 10-days | 100-days |
| M00307 | Pathway | Pyruvate oxidation | 2/6 | 1/5 | 1/6 |
| M00360 | FuncSet | Aminoacyl-tRNA biosynthesis | 3/6 | 1/5 | 3/6 |
| M00258 | Complex | Putative ABC transport system | 4/6 | 2/5 | 3/6 |
| M00049 | Pathway | Adenine ribonucleotide biosynthesis | 4/6 | 2/5 | 1/6 |
| M00083 | Pathway | Fatty acid biosynthesis | 3/6 | 2/5 | 3/6 |
| M00125 | Pathway | Riboflavin biosynthesis | 3/6 | 1/5 | 1/6 |
| M00015 | Pathway | Proline biosynthesis | 3/6 | 1/5 | 3/6 |
| M00048 | Pathway | Inosine monophosphate biosynthesis | 3/6 | 1/5 | 3/6 |
| M00120 | Pathway | Coenzyme A biosynthesis | 2/6 | 0/5 | 0/6 |
| M00240 | Complex | Iron complex transport system | 2/6 | 2/5 | 3/6 |

## Discussion

We found that the molecular evolutionary dynamics of phylogenetically distant taxa were consistently altered as energy limitation increased. In general, genetic diversity accumulated to a higher level than what we would expect if cellular division stopped after populations exited their exponential phase of growth after a single day. This consistent deviation suggests that some degree of cellular division continued while the population remained in an energy limited state, as it is doubtful that mutation alone would be capable of driving *de novo* mutations to detectable frequencies. While the biological mechanism responsible for this continued reproduction cannot be identified from mutation trajectory data, it is likely due to surviving cells scavenging dead cells, the only available resource, for energy^41,42^. This hypothesis is supported by the observation that the slope of *Bacillus* was closest to the null out of all taxa examined, which was likely due to *Bacillus* being the only taxon capable of forming non-reproductive endospores that buffer the accumulation of *de novo* genetic diversity^8,43^.

Comparisons of the molecular dynamics of different energy limitation regimes revealed that the frequency trajectories of mutations and the spectrum of nonsynonymous and synonymous polymorphic mutations *pN/pS* had higher similarity between 10 and 100-day regimes than either one did to the 1-day regime. A similar pattern was also found in our analyses of the direction of molecular evolution at the gene-level, where the degree of divergent evolution was at its lowest for comparisons made between 10 and 100-day regimes across taxa. This recurring pattern suggests that the main effect of energy limitation on molecular evolution occurred shortly after the populations exited their exponential phase of growth, an observation consistent with recent findings documenting the physiological and cellular changes that microorganisms incur after energy depletion^44^.

Despite the generality of the effect of energy limitation on the molecular evolutionary dynamics within a given taxon, the signal of this pattern dissipated when comparisons were made between phylogenetically distant taxa. By coarse-graining genes to a higher level of functional annotation, we found that convergent evolution occurred among phylogenetically distant taxa within certain energy limitation regimes. However, the same biological functions contributed to these signals of convergent evolution in energy repleted and depleted environments. Alternatively stated, while phylogenetically distant taxa acquired mutations at the same molecular targets within certain energy limitation regimes and divergent evolution occurred between energy limitation regimes within each taxon, the statistical signal of divergent evolution was lost when comparisons were made among all taxa. This lack of signal suggests that while energy limitation can consistently alter the rate and direction of molecular evolution in predictable ways, the targets of adaptation likely vary across phylogenetically distant taxa.

The lack of biological functions enriched for mutations within a specific energy limitation regime casts doubt on whether universal contributors towards adaptation can be identified by comparing evolve-and-resequence experiments from phylogenetically distant taxa. The absence of such contributors contrasts with the consistent effect that energy limitation has on evolutionary dynamics within a given taxon. This supposed contradiction can be resolved by accounting for the observation that energy limitation effects birth-death dynamics in a consistent manner across the microbial tree of life^42^, while proposing that different cellular mechanisms contribute towards adaptation within a given taxon. To compensate for this lack of signal, we argue that future efforts to investigate adaptation in energy limited environments should focus on characterizing the cellular effects of mutations that reach high frequencies. This approach, redolent of the framework of evolutionary cell biology^45,46^, would circumnavigate the issue of annotation conservation and allow researchers to instead examine conserved cellular features that contribute to adaption across phylogenetically distant species.

## Methods

### Long-term evolution under energy limitation

Energy limitation was manipulated by extending the time between transfers for microbial populations. We performed our energy limited evolution experiments using the following microbial taxa: *Bacillus subtilis* NCIB 3610 (ASM205596v1), *Caulobacter crescentus* NA1000 (ASM2200v1), *Deinococcus radiodurans* BAA-816 (ASM856v1), *Janthinobacterium* sp. KBS0711 (ASM593795v2), *Pedobacter* sp. KBS0701 (ASM593864v2), *Pseudomonas* sp. KBS0710 (ASM593804v2). A single colony was isolated from each taxon and grown in 10 mL of PYE media with 0.2% glucose and 0.1% casamino acids (2 g bactopeptone, 1 g yeast extract, 0.3 g MgSO_4_ x 7 H_2_O, 2 g dextrose, 1 g casamino acids in 1 L E-Pure™ H_2_O with 50 mL of a 10 g/mL cycloheximide solution added after autoclaving to prevent fungal contamination) in a 50 mL Erlenmeyer flask. A cultured flask of each taxon was transferred once a day for two days before being split between 15 flasks. Five replicates of each taxon were transferred as 1 mL aliquots into 9 mL of medium every 1, 10, or 100 days for ∼1,000 days. All flasks were organized in a 25°C shaker at 250 RPM such that no two populations of the same taxon were adjacent. One and 10 day treatments were cryopreserved in 20% glycerol at -80°C on days 30, 60, and 100 in 100-day increments, while 100-day treatments were cryopreserved every 100 days. Every 100 days biomass was cryopreserved for sequencing and each line was plated to examine contamination via CFU counts. Given that log_2_(10) ≈3.3 generations per-transfer, over a maximum of 1,000 days this experiment amounted to an evolutionary timescale of 3,300, 330, and 33 generations for 1-day, 10-day, and 100-day transfers, respectively. While we tested all taxa to make sure that they could survive 10 and 100 days without media replenishment, two 100-day *Pedobacter* populations and four 10-day *Janthinobacterium* populations repeatedly went extinct starting at day 400. Because of this reduction in temporal resolution we excluded 10-day *Janthinobacterium* populations from our analyses.

#### Signals of historical contingency in the distribution of extinction events

We compared, the coefficient of variation (CV) calculated from the observed distribution of extinction events to its null. Under a Poisson distribution, the coefficient of variation is 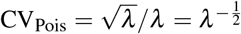 and observed values of CV greater than this null indicate that extinction events are overdispersed. Using the mean number of extinction events 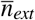 as an estimate of *λ* for all taxon-treatment combinations where at least three replicate populations went extinct, we generated null Poisson distributed extinction events using `numpy v1.16.4.` Multiple testing correction was performed for the *P*-values using the Benjamini–Hochberg procedure at *α* = 0.05

### Phylogenetic reconstruction

The 16S rRNA gene of each taxon was amplified via PCR using 8F and 1492R primers and reaction conditions previously described^47^. PCR products were purified using the QIAGEN QIAquick PCR Purification Kit and Sanger sequenced at the Indiana Molecular Biology Institute (IMBI) at Indiana University Bloomington (IUB). Sequences were aligned using SILVA INcremental Aligner v1.2.11 (`SINA`)^48^. Phylogenetic reconstruction was performed using Randomized Axelerated Maximum Likelihood v8.2.11 (`RAxML`)^49^ with the General Time Reversible model, gamma distributed rate variation, and bootstrap convergence criteria set to `autoMRE`. The 16S rRNA sequence of *Prochlorococcus marinus* subsp. marinus str. CCMP1375 (NC_005042) was used as an outgroup. Phylogenetic distance was calculated using the ETE Toolkit^50^.

### Library construction and sequencing

Cryopreserved biomass samples were mixed with 1 mL of a 50 mg/mL lysozyme (Dot Scientific DSL38100-10) E-Pure™ H_2_O solution and incubated at 37°C for 60 min. DNA extractions were performed using Qiagen DNeasy UltraClean Microbial Kits. Libraries were constructed out of 1,337 samples. Approximately 30% of Illumina libraries were constructed in-house with the Nextera DNA Library Preparation Kit (Illumina, FC-121-1030) using a protocol based off of previously published protocols^51,52^. These libraries were sequenced as paired-end reads on an Illumina HiSeq 2500 at the University of New Hampshire Hubbard Center for Genome Studies. All remaining samples were constructed as Illumina Nextera DNA paired-end 2 x 150bp libraries by the Center for Genomics and Bioinformatics at Indiana University and sequenced on a Illumina NextSeq 300. A target depth of 100-fold coverage was set for all libraries.

### Variant calling and analysis

The first 20 bp of all reads were trimmed and all read pairs where at least one pair had a mean Phred quality less than 20 were removed `cutadept v1.9.1`^53^. Candidate variants were identified using a previously published approach^33^. We provide a brief summary of the method in the supplement, including a high-level overview of the procedure, modifications made to accommodate our experimental design, and key parameter settings. To estimate mutation frequency for all significant candidates, we use the naive estimator 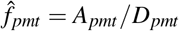. We examined the accumulation of mutations by time *t* as the sum of derived allele frequencies 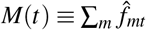 for each population. The change in *M*(*t*) between two timepoints was defined as Δ*M*(*t*) ≡ log_10_ [*M*(*t*)*/M*(*t*−1)]. The maximum observed frequency of a given mutation over *T* observations was defined as *f*_*max*_ ≡max({*f* (*t*): *t* = 1, …, *T*}), where *f* (*t*) is the mutation frequency at time *t*. Nucleotide transitions for all six spectra were counted at all mutations that occurred at four-fold redundant sites. The ratio of nonsynonymous to synonymous polymorphic mutations *pN/pS* was corrected by the number of nonsynonymous and synonymous sites within a genome. We examined *pN/pS* for values of *f*_*max*_ that were found across all taxa and treatments.

Principal component analysis was performed on the *n* × *p* matrix **X** containing *p* relativized nucleotide transitions for all *n* populations using `scikit-learn v0.23.1`^54^. Significant factor loadings were determined by creating permutation arrays *π* = (*π*_1_, *π*_2_, …, *π*_*p*_), where *π*_*j*_ is a permutation of the *n* populations for the *j*th transition. This permutation array generates the permuted data matrix **X**_*π*_, on which PCA was performed and null factor loadings were calculated. We performed 10,000 iterations and *P*-values were calculated by comparing observed factor loadings to their null distribution. A permutational multivariate ANOVA (PERMANOVA) was performed using a modified pseudo-*F* statistic to account for unequal covariance^55^.

### Within-taxon parallel and convergent/divergent evolution

For the purpose of this analysis, genetic parallelism is defined as the degree that the distribution of observed nonsynonymous mutation counts across genes deviates from the null expectation account based on gene size (i.e., multiplicity). Genes with significant multiplicity were identified using a previously published appr<u>o</u>ach^33^. To briefly summarize, we begin by calculating the multiplicity of a gene of size *L*_*i*_ with *n*_*i*_ mutations of size as 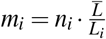, where 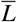 is the mean gene size in the genome. Under our null hypothesis, all genes have the same multiplicity 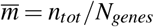. We then quantify the net increase of the log-likelihood of the alternative hypothesis relative to the null across the genome as 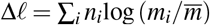. Additional details regarding this approach and how significant genes are identified can be found in its original publication^33^ and in the supplement.

Divergence and convergence as the presence or absence of enriched genes across energy limitation regimes within a given taxon was tested by determining whether the size of the intersection of the set of genes with significant multiplicity across treatments within a taxon was greater than (convergence) or less than (divergence) expected by chance. This can be performed by extending the hypergeometric distribution to the multivariate case, where the probability that *m* treatments have *k* intersecting genes is

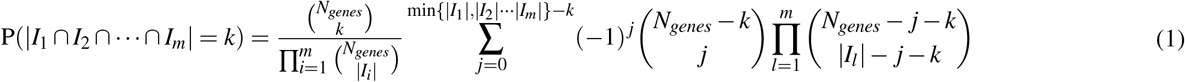

where *I*_*l*_ represents the set of genes that are significantly enriched for nonsynonymous mutations in the *l*th treatment and represents its cardinality. However, given that *N*_*genes*_∼*O*(10^3^), it can be prohibitive to calculate and multiply all binomial coefficients inside the sum and the fact that the cardinality varied across treatments prevented us from approximating the sum as a Gaussian distribution. Instead we simulated the process described by Eq.1 to determine whether the Jaccard similarity of gene content between two treatments (*J*(*I*_*i*_, *I*_*j*_) = *I*_*i*_∩*I*_*j*_ */ I*_*i*_∪*I*_*j*_) was greater or less than expected by chance.

Building off of prior results^56^, the degree of divergent versus convergent evolution among enriched genes was determined using the squared Pearson’s correlation of relative multiplicities (*µ*_*i*_ = *m*_*i*_*/* ∑ *m*_*i*_) for each pair of energy limitation regimes. Null correlation coefficients were calculated by constructing a gene-by-regime mutation count matrix for each pair of energy limitation regimes within each taxon and randomizing combinations of mutation counts constrained on the total number of mutations acquired within each gene across treatments and the number of mutations acquired within each treatment (i.e., the row and column sums of the matrix). This randomization was performed using a Python implementation^57^ of the ASA159 algorithm^58^. Correlation coefficients were calculated from the randomized matrices 10,000 times and the observed coefficients were standardized relative to their respective null distributions (*Z*_*ρ*_).

### Among-taxa convergent evolution

Because few genes are present among all of the taxa we chose, it is difficult to determine whether the same gene acquired more mutations than expected by chance across multiple taxa (i.e., evolutionary convergence among taxa within the same energy limitation regime). To determine whether evolutionarily distant taxa acquired similar mutations, we grouped genes at higher levels of biological organization. To do this, we first inferred the metabolic pathway composition of all taxa using the Metabolic And Physiological potentiaL Evaluator (`MAPLE v2.3.1`;^39^). `MAPLE` was run using bi-directional best hit with NCBI `BLAST` on KEGG genes and modules version `20190318` using all bacteria and archaea in the database. MAPLE output files for the module pathways, signatures, and complexes present in all six taxa were filtered for query coverage values of 100% and merged into a single file for each taxon. A map was created using RefSeq protein IDs and KEGG annotations for each taxon and KEGG annotations were obtained for each gene with significant multiplicity. These KEGG annotations were mapped to the sub-set of MAPLE modules that passed our filtering criteria, where we finally generate a MAPLE module-by-taxon presence absence matrix. PERMANOVA was performed using the same approach, but with a modified permutation matrix that only permuted treatment labels within a given taxon.

Mapping the enriched genes to a higher level of biological organization required our null statistical model to be more complex than what was described in Eq.1. First, not all genes in all genomes are able to be annotated by KEGG. Second, KEGG annotated genes can be part of multiple MAPLE modules. That is, the relationship between the set of KEGG genes and MAPLE modules is not, strictly speaking, a mathematical function. However, because we are only interested in the degree of overlap in MAPLE modules across taxa within a given treatment we do not need to define the map between KEGG and MAPLE annotations. Rather, we can generate a null distribution for the size of intersecting sets of MAPLE modules by randomly sampling KEGG genes, mapping them to MAPLE pathways, and generating null MAPLE pathway-by-taxon matrices. Because each taxon has a distinct annotation, we set the number of significant genes that also had KEGG annotations as the sample size. Significance of a given intersection set size was established by comparing observed data to the null distributions, where for set intersections with multiple combinations we summed over the 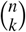 possible combinations for *k* intersections among *n* total taxa. Significance of the relationship between phylogenetic distance and the Jaccard index of MAPLE modules was assessed by comparing the slope obtained by ordinary least squares regression to a null distribution generated by calculating the OLS slope of the permuted variables for 10,000 iterations. Slope significance of distance-decay relationships was done as previously described^59^, where the absolute difference in slope values between a given pair of transfer regimes 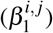 was compared to a null distribution generated by randomly permuting treatment labels and calculating the difference in slopes.

## Data availability

Raw sequence data is available on the NCBI Sequencing Read Archive under BioProject ID PRJNA639414. Reproducible code to perform the analyses in this study is available on GitHub under the repository: Phylo_Evol_Timeseries. Processed data and annotations are available on Zenodo under the DOI: 10.5281/zenodo.4517573.

## Acknowledgements

We thank J. Weissman and R. Wolff for providing valuable feedback on an earlier version of this manuscript. We thank M. Behringer, T. Doak, D. A. Drummond, P. Foster, M. Lynch, J. McKinlay, and members of the Lennon lab for their insights at various stages of the project. We thank H. Long for his help in setting up library construction and K. Miller for assisting with sample collection. This work was supported by US Army Research Office Grant W911NF-14-1-0411. Computing resources for simulations was supported by Lilly Endowment, Inc., through its support for the Indiana University Pervasive Technology Institute, the National Science Foundation under Grant No. CNS-0521433, and Shared University Research grants from IBM, Inc., to Indiana University.

## Author contributions

W.R.S., E.P., M.H.G., and J.T.L. designed the project; W.R.S., E.P., and M.H.G. conducted the experiments and generated the data; W.R.S. performed all statistical analyses; W.R.S., E.P., M.H.G., and J.T.L. wrote the manuscript.

