## Supplemental Information for "Molecular evolutionary dynamics of energy limited microorganisms"

#### 1 Variant calling

The first 20 bp of all reads were trimmed and all read pairs where at least one pair had a mean Phred quality less than 20 were removed `cutadept v1.9.1` [1]. Candidate variants were identified by modifying the codebase of a previously published approach under GPL v2 [2]. We provide a brief overview of the method used in Good et al. (2017) and all alterations made, though we direct the reader to the original publication for an in-depth account of the pipeline. Using `breseq v0.32.0` [3] to align trimmed reads to the reference genome, we generated a list of candidate junctions for all samples. Candidate junctions were merged across temporal samples for each taxon using `gdtools`, which was then passed as an argument to a second round of `breseq` using `--user-evidence-gd`. Using `samtools mpileup` [4] and open access Python scripts from [2], we identified candidate SNVs and small indels from BAM files generated by the second `breseq` run. We defined trajectories of ordered pairs ($A_{pmt}$, $D_{pmt}$) for the alternative allele count and total depth of coverage for mutation $m$ in sample $t$ from population $p$. Alternative alleles for indels $< 100$bp are merged as a single "compound" mutation trajectory. Indels $\geq 100$bp are considered structural variants and are processed with candidate junctions identified by `breseq`.

Candidate junctions were processed using a custom Python script that merges similar candidate junctions from the same population into a single "compound" junction candidate. Each of these candidates was recorded as a single trajectory ($A_{pmt}$, $D_{pmt}$). We examined all candidate mutations with $A_{pmt} \geq 2$ in at least two samples and $D_{pmt} \geq 10$ in at least one samples, with an observed frequency $f_{pmt} \equiv A_{pmt}/D_{pmt} \geq 0.05$ .

Statistical support of each candidate mutation was established using two summary statistics: 1) the autocorrelation of frequencies between timepoints ($C^*$) and 2) the derived allele sojourn weight ($I$). Both functions are described at length in Good et al. (2017). For our purposes here $C^*$ can be described as a modified form of the autocorrelation function

$$C \equiv \sum_t \left(f_{t+1} - \overline{f}\right)\left(f_t - \overline{f}\right) \quad (1)$$

where $f_t$ is the frequency of the mutation at a given timepoint and $\overline{f}$ is the mean frequency over the entire timecourse. We use a modified form of this function that has the same purpose, but accounts for discreteness and uncertainty in $f_t$ due to finite coverage.

The derived allele sojourn time is less intuitive. $C^*$ treats positive and negative deviations from the mean the same way (i.e., symmetric). Real mutations can have low frequencies for extended periods of time. A large area under this allele frequency trajectory curve would suggest that this mutation is not an error. To capture this trend, the statistic in Good *et al.* (2017) examines runs of 2 or more timepoints where $f_t$ is larger than a threshold frequency $f^*$ for that entire run. The run with the largest value of

$$I = \sum_{t=t_1}^{t_2} = f_t - f^* \tag{2}$$

The value of $f^*$ was chosen using the criteria in Good *et al.* to be as low as possible while allowing for error rates higher than $1/D_t$ (2017).

The two $P$-values were merged as a composite $P$-value using the following function:

$$T = \sum_k \theta\left(P^* - P_k\right) \log\left(\frac{1}{P_k}\right) \tag{3}$$

where $\theta(\cdot)$ is the Heaviside step function and $P^* = (0.05)^{1/2}$. Significance was assessed by calculating the null distribution of $T$ for all mutations and nonsignificant mutations were removed from downstream analyses. We did not include the third statistic, the average frequency relaxation time, used in Good *et al.* due to fact that our experiment covers a relatively brief evolutionary timescale ( 3,000 generations vs. 60,000) (2017). We annotated all mutations as described in Good *et al.* (2017).

### 2 Mutation trajectory inference

We use the naive estimator $\hat{f}_{pmt} = A_{pmt}/D_{pmt}$ as our measure of mutation frequency. We examined mutation accumulation as

$$M(t) \equiv \sum_{m} \hat{f}_{pmt} \tag{4}$$

a measure that uses information from mutation frequencies in addition to the number of mutations in the population.

To infer whether a high frequency mutation is present in all individuals (i.e., "fixed"), we used the hidden Markov model introduced in Good *et al.* with ($A_{mt}$;$D_{mt}$) as the observed sequence of emissions (2017). To briefly summarize this model, mutations start in an ancestral state **A** where $f_{mt} = 0$. At each timepoint mutations can transition to a polymorphic state **P** where the frequency remains as $0 < f_{mt} < 1$. From here the mutation can either transition to a fixed state **F** or go extinct **E**. **E** states are allowed to re-appear as **A**, though multiple mutations occurring at the same site is a rare event. The behavior of the HMM was fairly insensitive to the chosen initial transition probabilities, so we set the initial transition probabilities to those in Good *et al.* (2017).

### 3 Parallelism and divergence

We identified potential targets of selection by examining the distribution of nonsynonymous mutations across genes. The statistical framework of this approach was developed in Good *et al.* (Good et al., 2017). To briefly summarize, gene-level parallelism was assessed by calculating the *multiplicity* of each gene as

$$m_i = n_i \cdot \frac{\overline{L}}{L_i} \tag{5}$$

where $n_i$ and $L_i$ is the number of mutations observed and the length of the $i$th gene and $\overline{L}$ is the mean length of all genes. Under this definition, the null hypothesis is that all genes have the same multiplicity $\overline{m} = n_{tot}/N_{genes}$. Using the observed and expected values, we can quantify the net increase of the log-likelihood of the alternative hypothesis relative to the null

$$\Delta\ell = \sum_i n_i \log\left(\frac{m_i}{\overline{m}}\right) \tag{6}$$

where significance is assessed using permutation tests. Because this measure can be sensitive to $n_{tot}$, for comparisons across different strains and treatments we randomly sub-sampled mutations as a multinomial distribution, where the probability of sampling a mutation at gene $i$ was given by $p_i = n_i/n_{tot}$. Multinomial sampling was performed 10,000 times with a sub-sampled $n_{tot}$ set to 50.

To identify specific genes that are enriched for mutations, we calculated the $P$-value of each gene as

$$P_i = \sum_{n \geq n_i} \frac{\left(\frac{n_{tot} L_i}{\overline{L} N_{genes}}\right)^n}{n!} e^{-\frac{n_{tot} L_i}{\overline{L} N_{genes}}} \tag{7}$$

where FDR correction was performed by defining a critical $P$-value ($P^*$) based on the survival curve of a Poisson distribution, the null hypothesis (see Good et al., 2017 for additional details). We then defined the set of significant genes for each treatment-strain combination as:

$$I = \{i : P_i \leq P^*(\alpha)\} \tag{8}$$

for $\alpha = 0.05$.

| PC | Transition | $\rho^2$ | $P$ |
|---|---|---|---|
| 1 | A : T $\rightarrow$ C : G | 0.906 | $< 10^{-4}$ |
| | G : C $\rightarrow$ T : A | 0.746 | $< 10^{-4}$ |
| | G : C $\rightarrow$ C : G | 0.627 | 0.015 |
| 2 | A : T $\rightarrow$ G : C | 0.758 | 0.002 |
| | G : C $\rightarrow$ A : T | 0.657 | 0.015 |

Table S1: Factor loadings that are significantly correlated with the first and second components of the PCA visualized in Fig. 2.

| Genus | Treatment intersection | Size | $P$ |
|---|---|---|---|
| *Bacillus* | $1 \cap 10$ | 34 | $< 10^{-3}$ |
| | $1 \cap 100$ | 37 | $< 10^{-3}$ |
| | $10 \cap 100$ | 42 | $< 10^{-3}$ |
| | $1 \cap 10 \cap 100$ | 31 | $< 10^{-3}$ |
| *Caulobacter* | $1 \cap 10$ | 134 | $< 10^{-3}$ |
| | $1 \cap 100$ | 145 | $< 10^{-3}$ |
| | $10 \cap 100$ | 173 | $< 10^{-3}$ |
| | $1 \cap 10 \cap 100$ | 112 | $< 10^{-3}$ |
| *Deinococcus* | $1 \cap 10$ | 62 | $< 10^{-3}$ |
| | $1 \cap 100$ | 63 | $< 10^{-3}$ |
| | $10 \cap 100$ | 61 | $< 10^{-3}$ |
| | $1 \cap 10 \cap 100$ | 47 | $< 10^{-3}$ |
| *Pedobacter* | $1 \cap 10$ | 26 | $< 10^{-3}$ |
| | $1 \cap 100$ | 18 | $< 10^{-3}$ |
| | $10 \cap 100$ | 21 | $< 10^{-3}$ |
| | $1 \cap 10 \cap 100$ | 16 | $< 10^{-3}$ |
| *Janthinobacterium* | $1 \cap 100$ | 72 | $< 10^{-3}$ |
| *Pseudomonas* | $1 \cap 10$ | 76 | $< 10^{-3}$ |
| | $1 \cap 100$ | 136 | $< 10^{-3}$ |
| | $10 \cap 100$ | 135 | $< 10^{-3}$ |
| | $1 \cap 10 \cap 100$ | 70 | $< 10^{-3}$ |

Table S2: The number of genes that are enriched for a given treatment intersection. $P$-values correspond to tests of whether a given intersection size is greater than expected by chance.

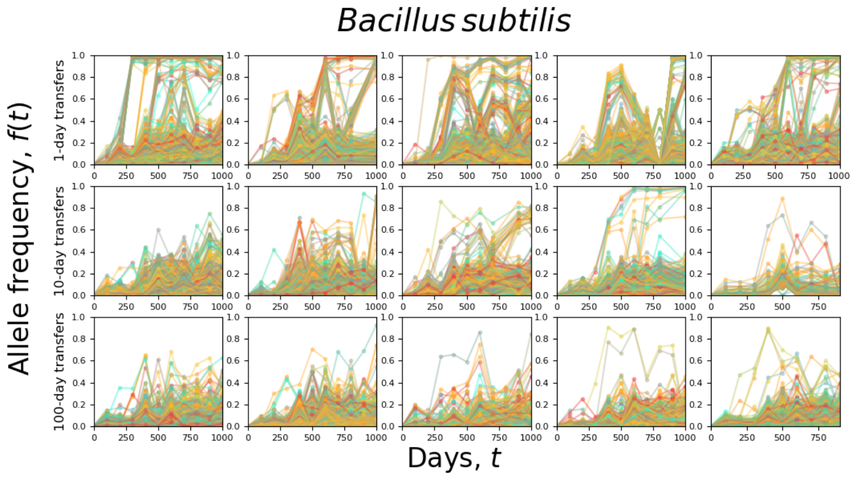


Figure S1: Allele frequency trajectories of all *Bacillus subtilis* replicate populations.

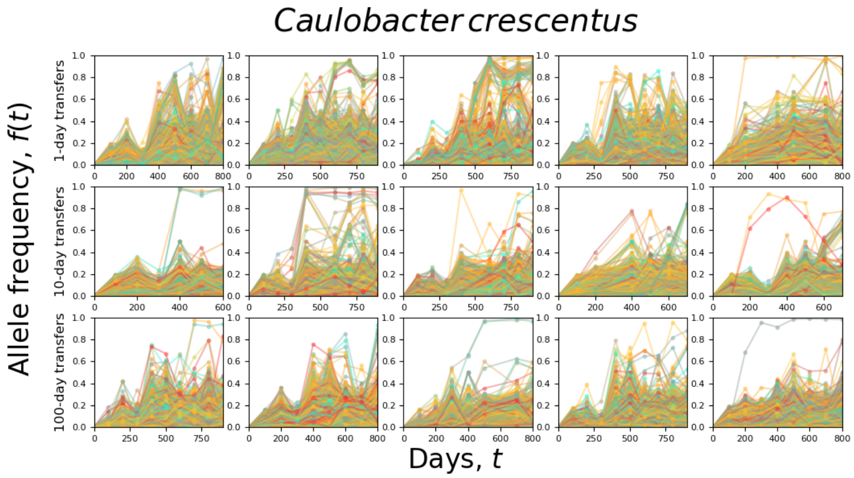


Figure S2: Allele frequency trajectories of all *Caulobacter crescentus* replicate populations.

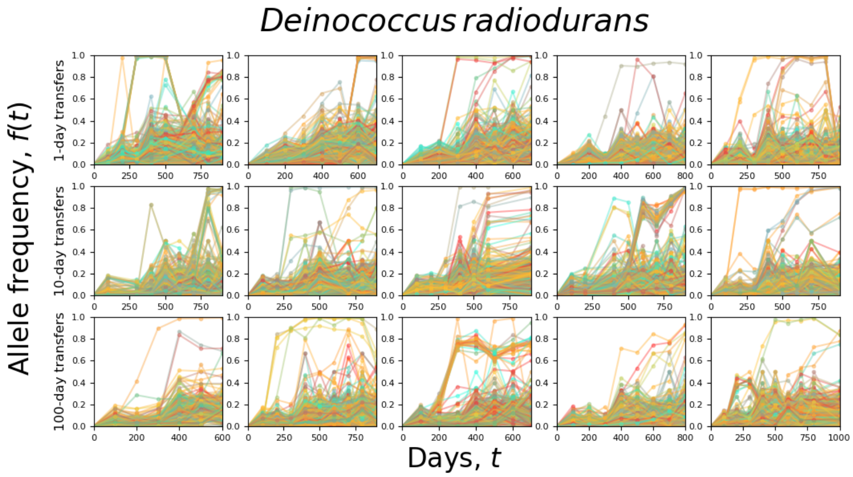


Figure S3: Allele frequency trajectories of all *Deinococcus radiodurans* replicate populations.

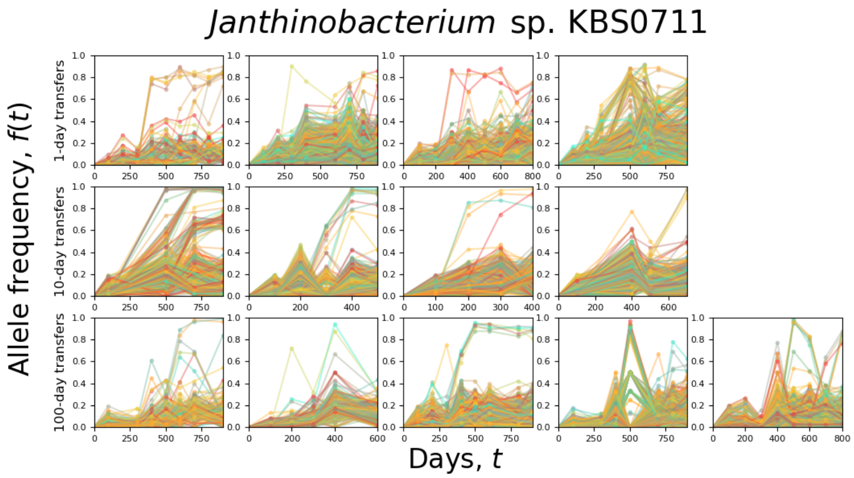


Figure S4: Allele frequency trajectories of all *Janthinobacterium* sp. KBS0711 replicate populations. Three 100-day replicate populations repeatedly went extinct over the course of the experiment, limiting the number of timepoints and preventing us from inferring mutation trajectories.

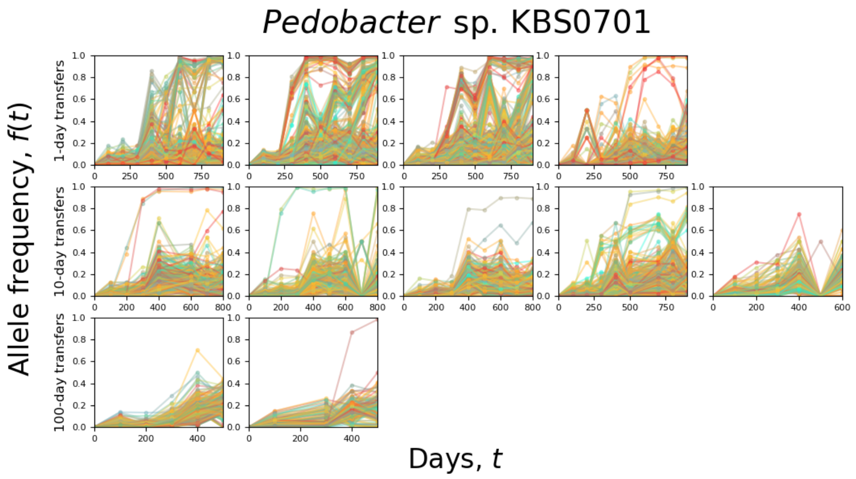


Figure S5: Allele frequency trajectories of all *Pedobacter* sp. KBS0701 replicate populations. Three 100-day replicate populations repeatedly went extinct over the course of the experiment, limiting the number of timepoints and preventing us from inferring mutation trajectories.

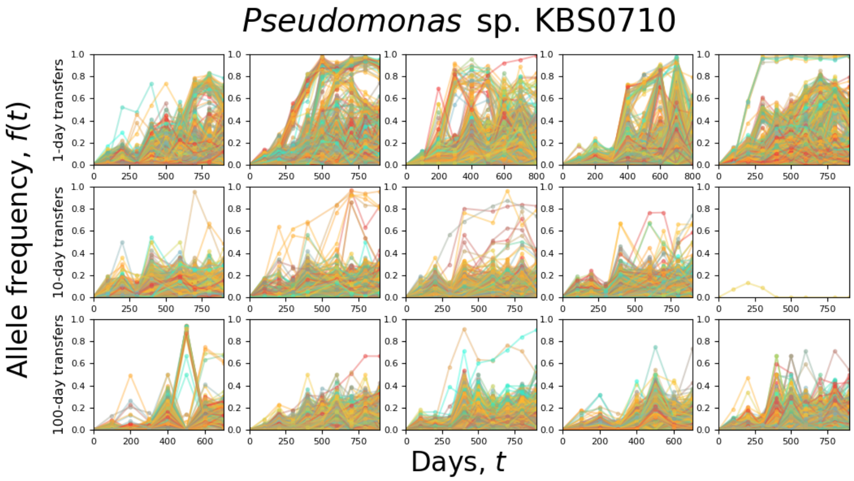


Figure S6: Allele frequency trajectories of all *Pseudomonas sp. KBS0710* replicate populations. Two 100-day replicate populations repeatedly went extinct over the course of the experiment, limiting the number of timepoints and preventing us from inferring mutation trajectories.

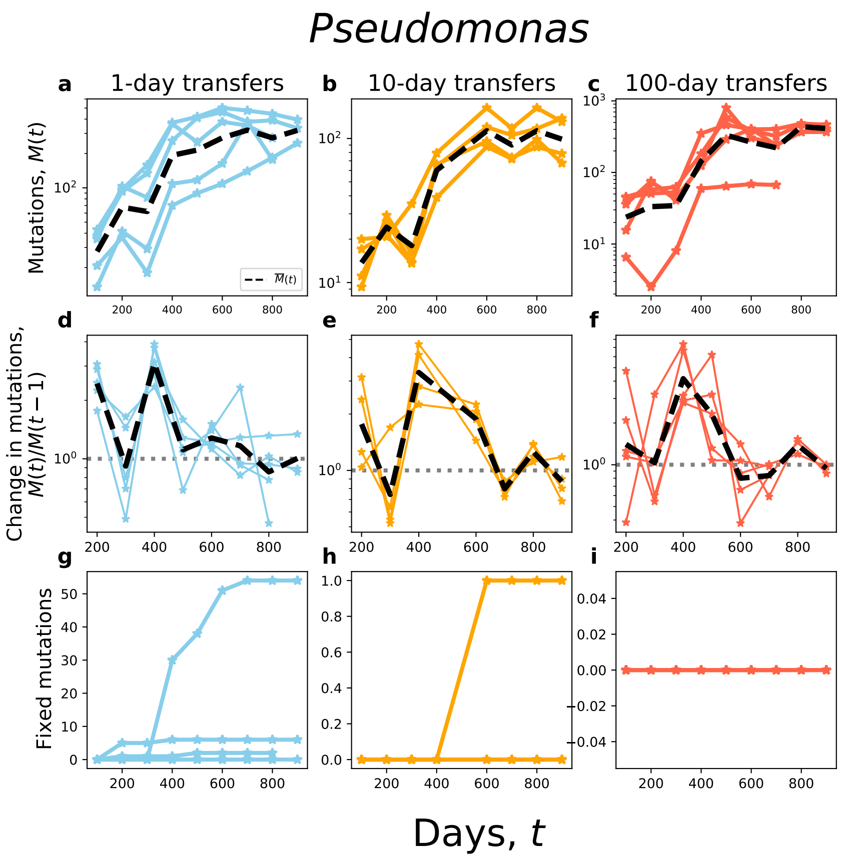


Figure S7: Molecular evolutionary dynamics of *Bacillus subtilis*. **a-c** Cumulative mutation ($M(t)$) trajectories for both strains over time. The dashed black line is the mean. **d-f** The change in $M(t)$ between timepoints across treatments. **g-i** The cumulative number of fixed mutations over time for all populations within a given treatment.

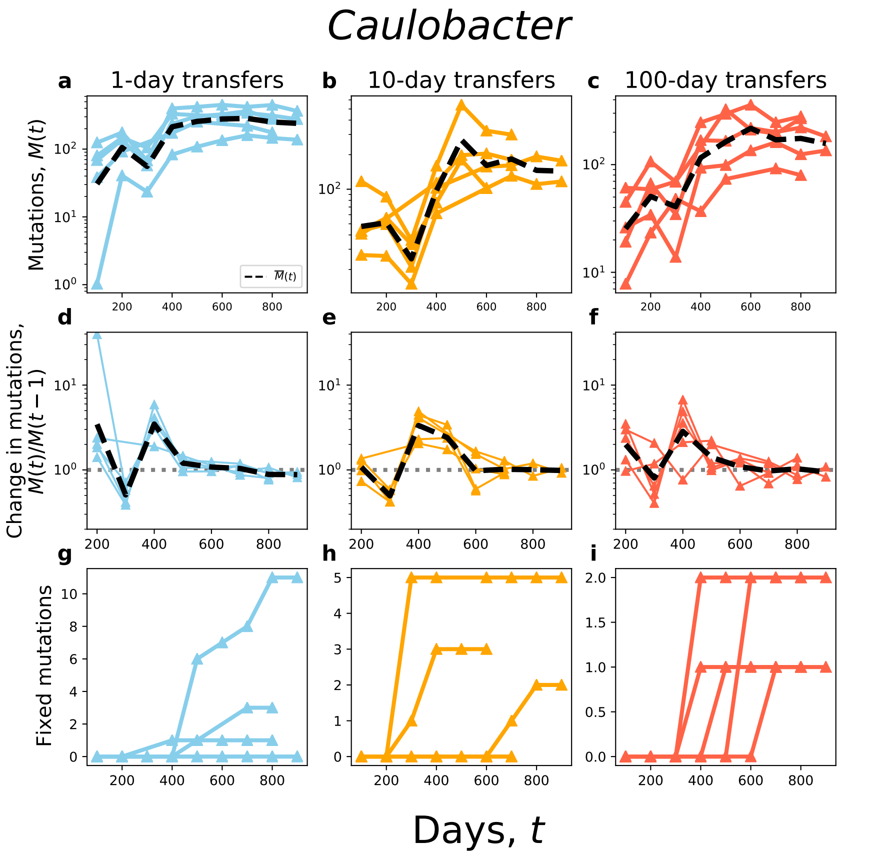


Figure S8: Molecular evolutionary dynamics of *Caulobacter crescentus*. **a-c** Cumulative mutation ($M(t)$) trajectories for both strains over time. The dashed black line is the mean. **d-f** The change in $M(t)$ between timepoints across treatments. **g-i** The cumulative number of fixed mutations over time for all populations within a given treatment.

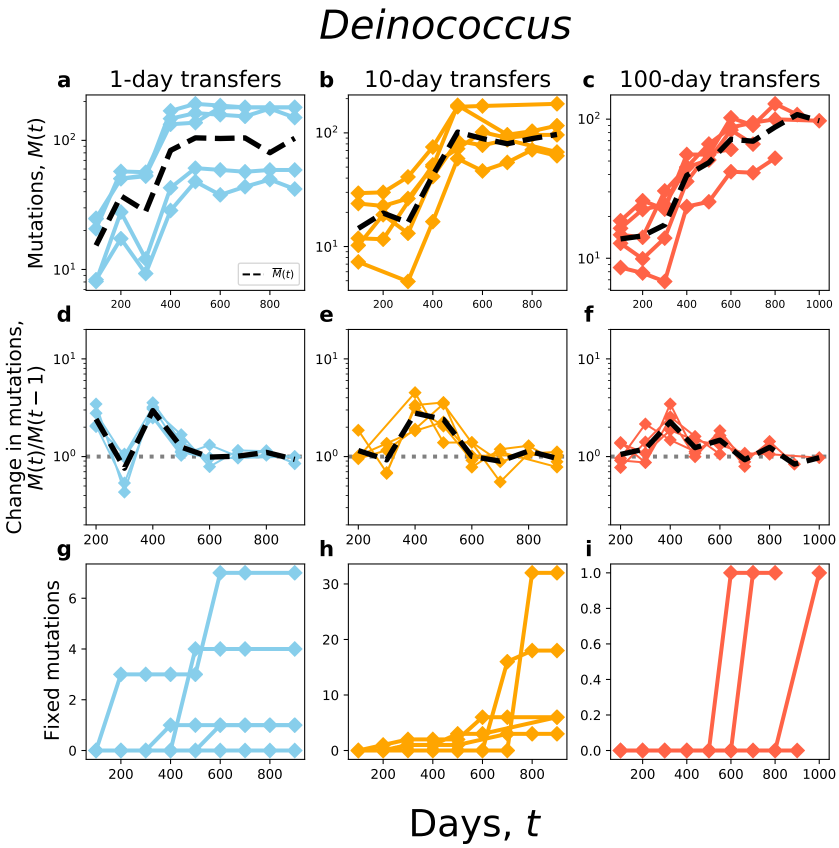


Figure S9: Molecular evolutionary dynamics of *Deinococcus radiodurans*. **a-c** Cumulative mutation ($M(t)$) trajectories for both strains over time. The dashed black line is the mean. **d-f** The change in $M(t)$ between timepoints across treatments. **g-i** The cumulative number of fixed mutations over time for all populations within a given treatment.

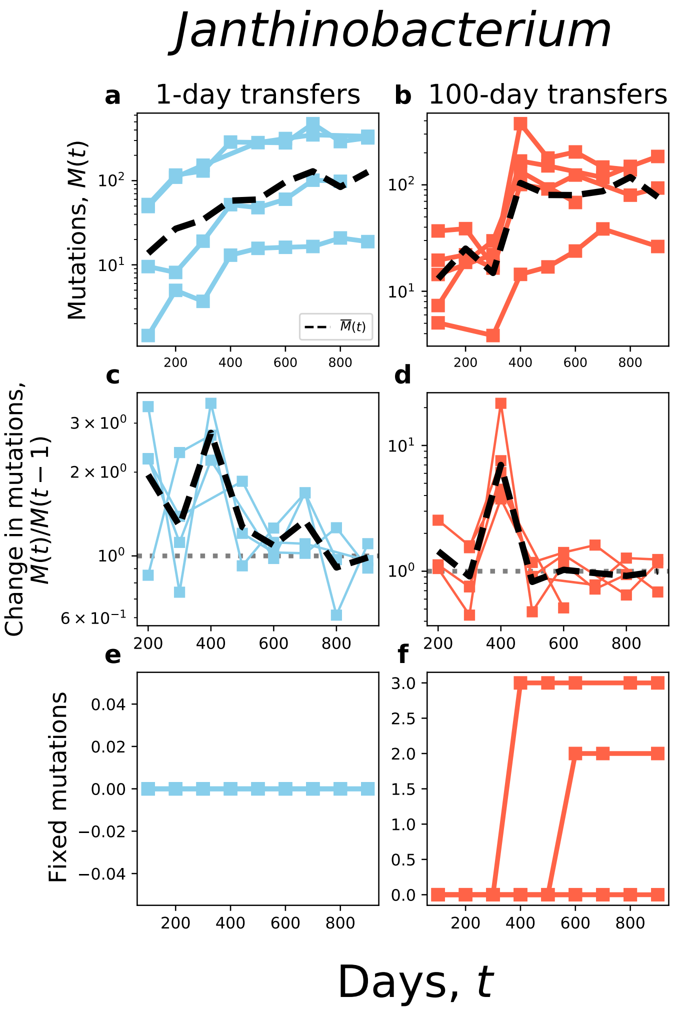


Figure S10: Molecular evolutionary dynamics of *Janthinobacterium* sp. KBS0711. **a-c** Cumulative mutation $(M(t))$ trajectories for both strains over time. The dashed black line is the mean. **d-f** The change in $M(t)$ between timepoints across treatments. **g-i** The cumulative number of fixed mutations over time for all populations within a given treatment.

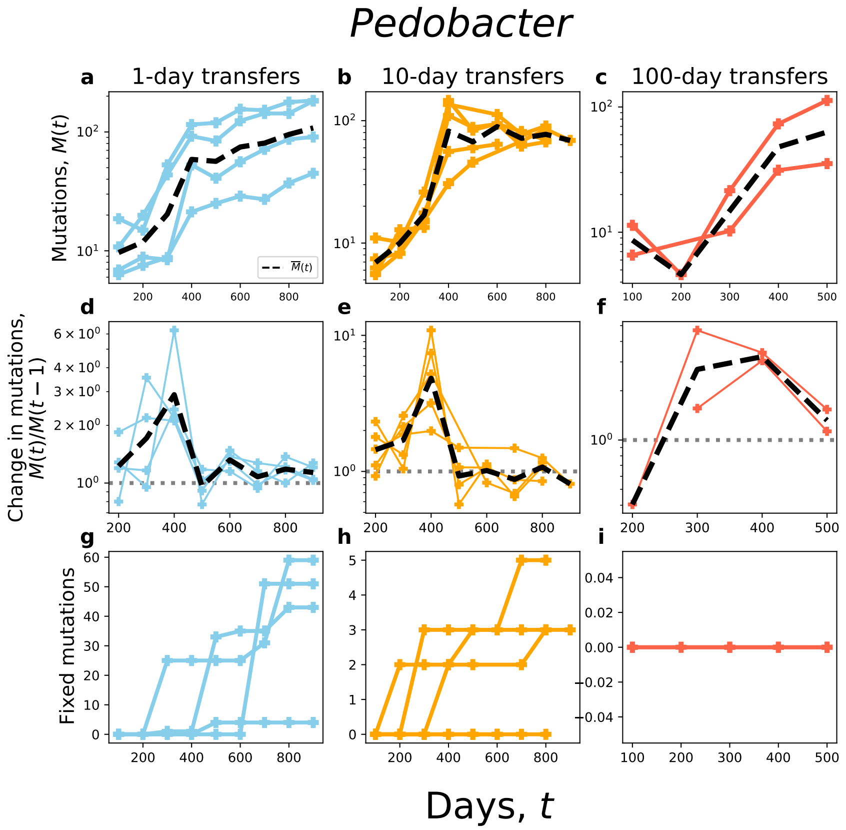


Figure S11: Molecular evolutionary dynamics of *Pedobacterium* sp. KBS0701. **a-c** Cumulative mutation ($M(t)$) trajectories for both strains over time. The dashed black line is the mean. **d-f** The change in $M(t)$ between timepoints across treatments. **g-i** The cumulative number of fixed mutations over time for all populations within a given treatment.

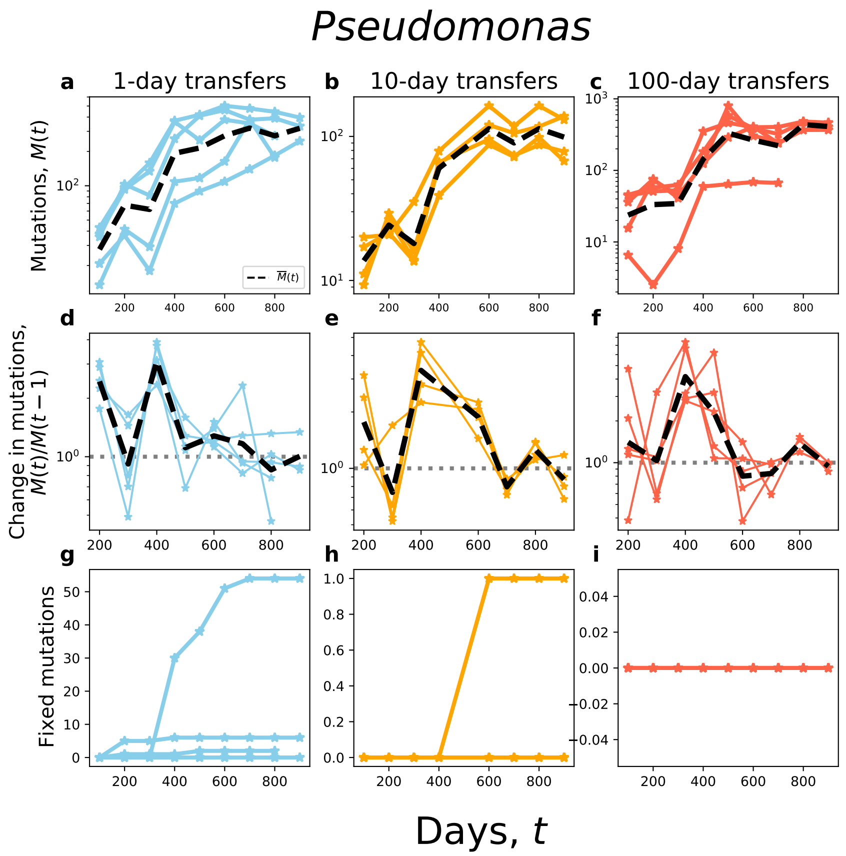


Figure S12: Molecular evolutionary dynamics of *Pseudomonas* sp. KBS0710. **a-c** Cumulative mutation ($M(t)$) trajectories for both strains over time. The dashed black line is the mean. **d-f** The change in $M(t)$ between timepoints across treatments. **g-i** The cumulative number of fixed mutations over time for all populations within a given treatment.

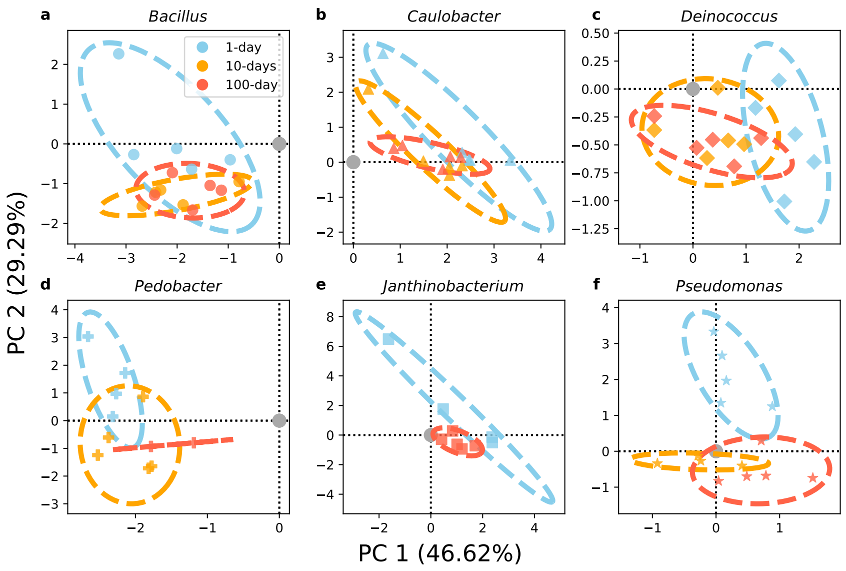


Figure S13: **a-f)** PCA of observed mutation spectra for all replicate populations, where each taxon has been plotted separately with 95% confidence ellipses for each treatment.

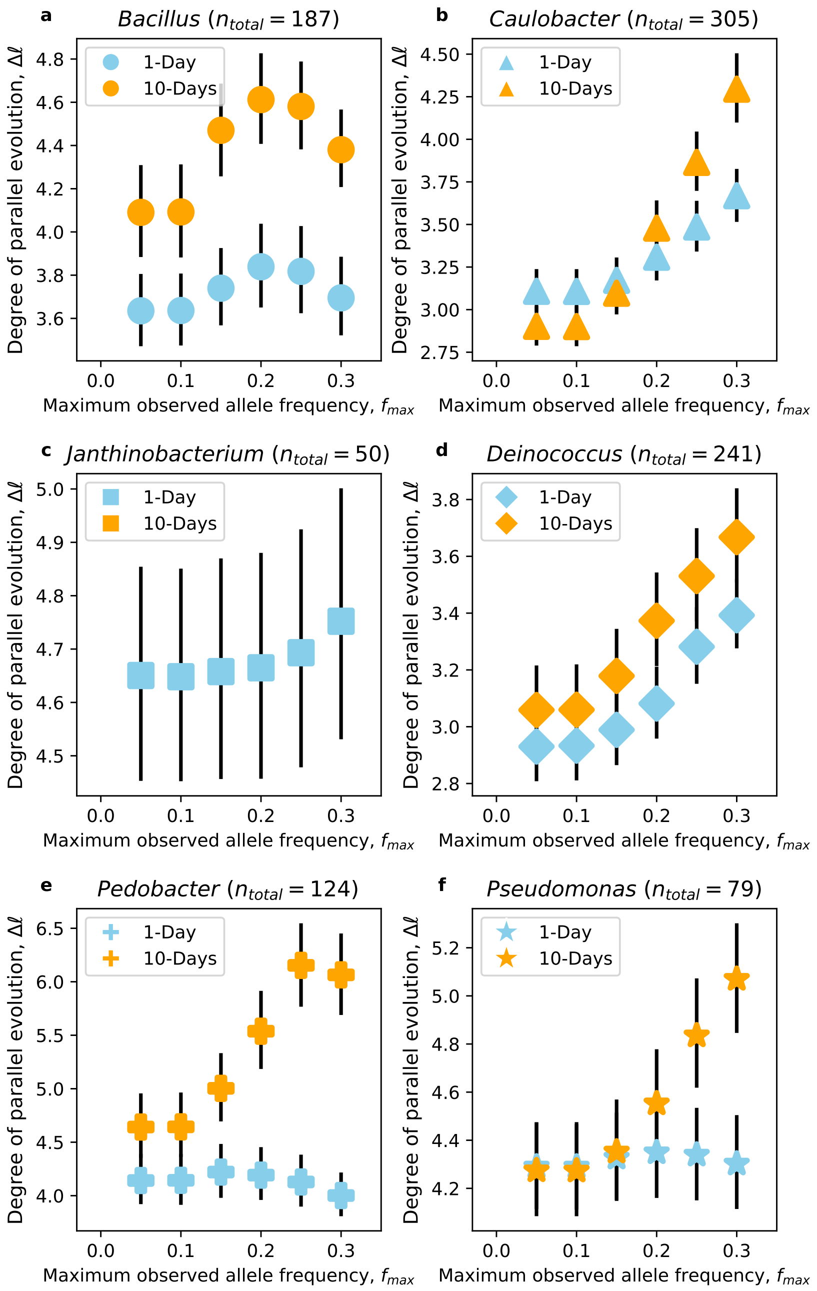


Figure S14: **a-f)** **a-f)** The relationship between the maximum observed frequency of a mutation ($f_{max}$) and the degree of parallelism. Genome-wide parallelism is generally higher among mutations that reach a higher maximum frequency during their sojourn time in the population. Dots and bars represent the mean and 95% CIs from 10,000 subsamples of mutations with a given $f_{max}$ cutoff, respectively.

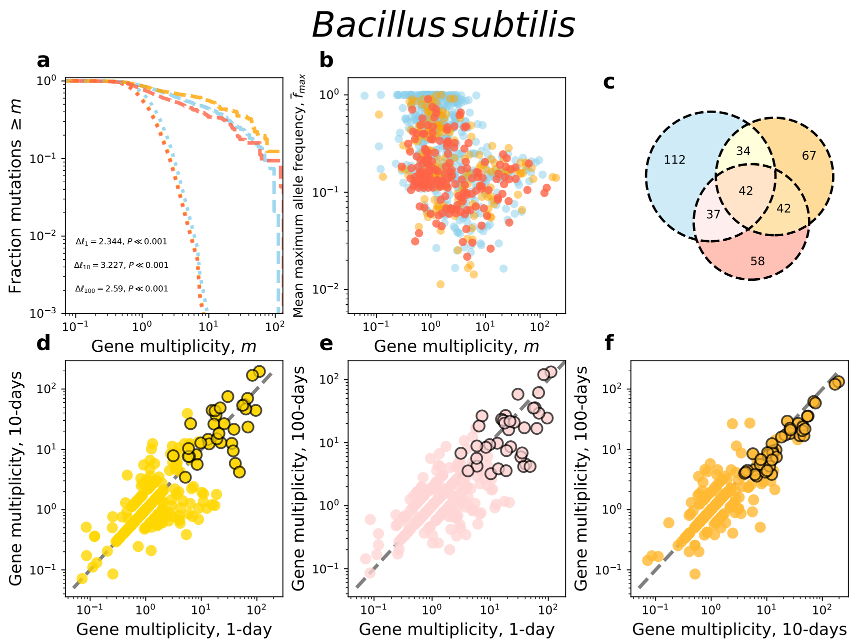


Figure S15: Parallelism and divergence/convergence visualizations for all mutations in *Bacillus*. **a**) The survival curve for multiplicity decays at a slower rate than the null across treatments, indicating that we can reject the null hypothesis that nonsynonymous mutations are equally distributed across genes. **b**) A scatterplot visualizing the relationship between the mean maximum frequency ($\overline{f}_{max}$) and multiplicity of genes. **c**) A venn diagram showing the overlap in genes that were significantly enriched for nonsynonymous mutations across treatments. **d-f**) Pairwise comparisons of multiplicity for genes across all treatments, where significantly enriched genes have a black outline.

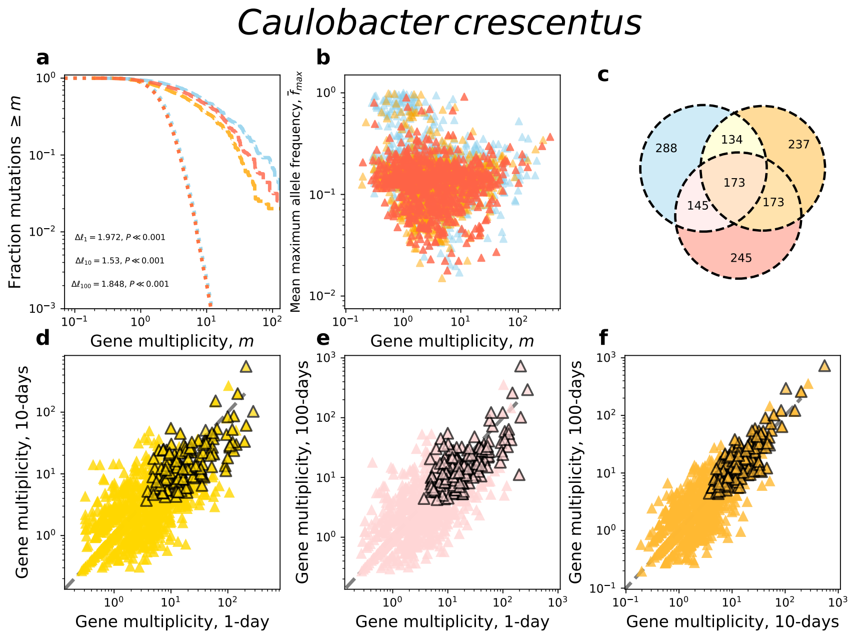


Figure S16: Parallelism and divergence/convergence analyses for all mutations in *Caulobacter*. **a**) The survival curve for multiplicity decays at a slower rate than the null across treatments, indicating that we can reject the null hypothesis that nonsynonymous mutations are equally distributed across genes. **b**) A scatterplot visualizing the relationship between the mean maximum frequency ($\overline{f}_{max}$) and multiplicity of genes. **c**) A venn diagram showing the overlap in genes that were significantly enriched for nonsynonymous mutations across treatments. **d-f**) Pairwise comparisons of multiplicity for genes across all treatments, where significantly enriched genes have a black outline.

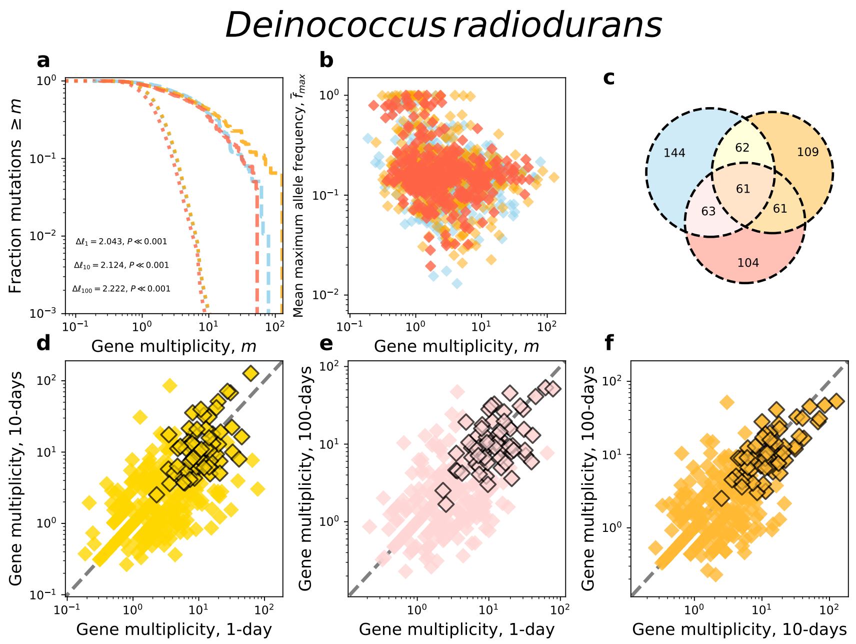


Figure S17: Parallelism and divergence/convergence analyses for all mutations in *Deinococcus*. **a**) The survival curve for multiplicity decays at a slower rate than the null across treatments, indicating that we can reject the null hypothesis that nonsynonymous mutations are equally distributed across genes. **b**) A scatterplot visualizing the relationship between the mean maximum frequency ($\overline{f}_{max}$) and multiplicity of genes. **c**) A venn diagram showing the overlap in genes that were significantly enriched for nonsynonymous mutations across treatments. **d-f**) Pairwise comparisons of multiplicity for genes across all treatments, where significantly enriched genes have a black outline.

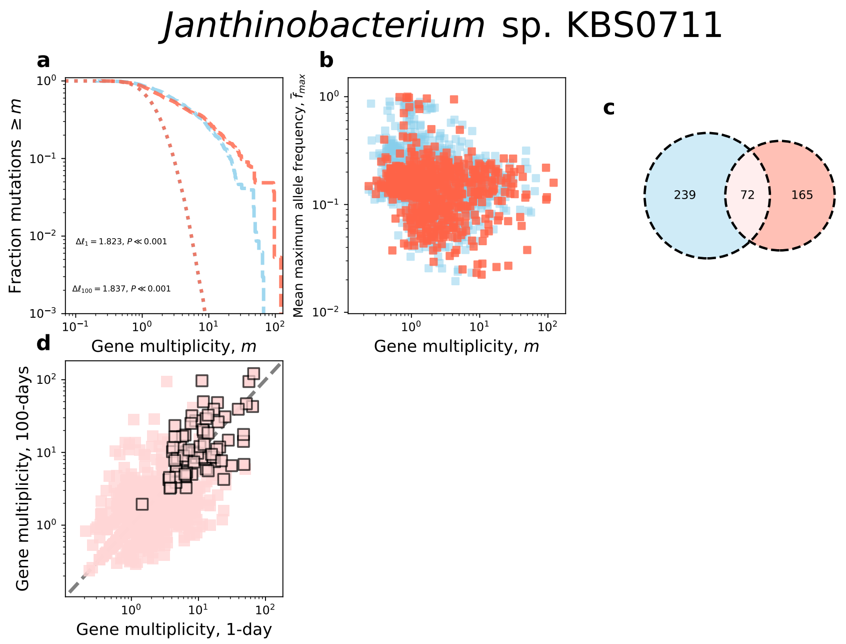


Figure S18: Parallelism and divergence/convergence analyses for all mutations in *Janthinobacterium*. **a**) The survival curve for multiplicity decays at a slower rate than the null across treatments, indicating that we can reject the null hypothesis that nonsynonymous mutations are equally distributed across genes. **b**) A scatterplot visualizing the relationship between the mean maximum frequency ($\overline{f}_{max}$) and multiplicity of genes. **c**) A venn diagram showing the overlap in genes that were significantly enriched for nonsynonymous mutations across treatments. **d-f**) Pairwise comparisons of multiplicity for genes across all treatments, where significantly enriched genes have a black outline.

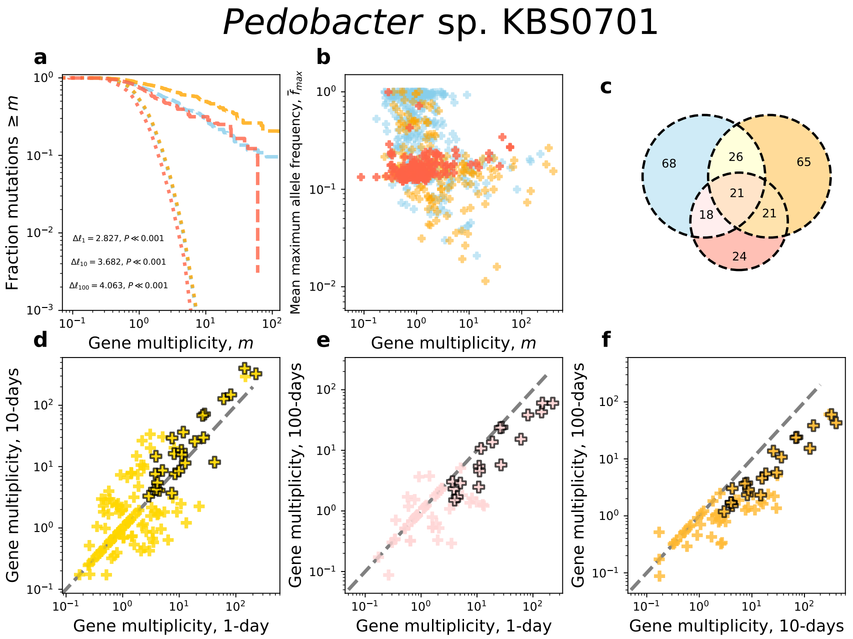


Figure S19: Parallelism and divergence/convergence analyses for all mutations in *Pedobacter*. **a**) The survival curve for multiplicity decays at a slower rate than the null across treatments, indicating that we can reject the null hypothesis that nonsynonymous mutations are equally distributed across genes. **b**) A scatterplot visualizing the relationship between the mean maximum frequency ($\overline{f}_{max}$) and multiplicity of genes. **c**) A venn diagram showing the overlap in genes that were significantly enriched for nonsynonymous mutations across treatments. **d-f**) Pairwise comparisons of multiplicity for genes across all treatments, where significantly enriched genes have a black outline.

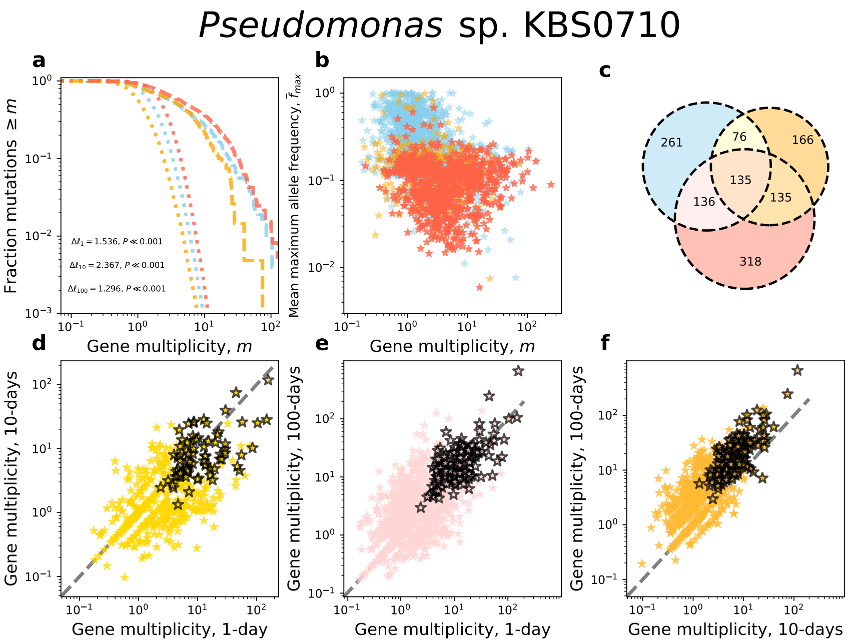


Figure S20: Parallelism and divergence/convergence analyses for all mutations in *Pseudomonas*. **a**) The survival curve for multiplicity decays at a slower rate than the null across treatments, indicating that we can reject the null hypothesis that nonsynonymous mutations are equally distributed across genes. **b**) A scatterplot visualizing the relationship between the mean maximum frequency ($\overline{f}_{max}$) and multiplicity of genes. **c**) A venn diagram showing the overlap in genes that were significantly enriched for nonsynonymous mutations across treatments. **d-f**) Pairwise comparisons of multiplicity for genes across all treatments, where significantly enriched genes have a black outline.

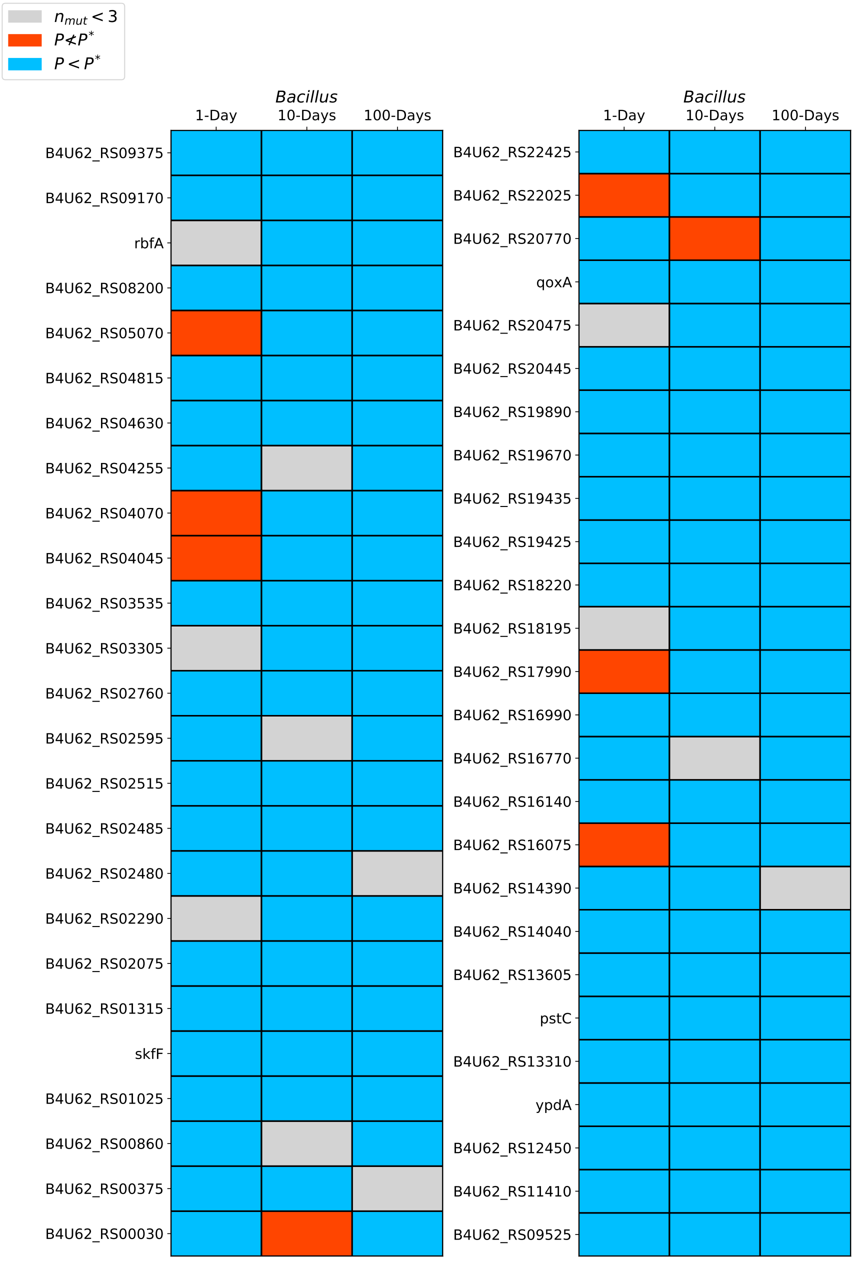


Figure S21: Visualization of all genes with an excess of non-synonymous mutations in more than one treatment for *Bacillus*.

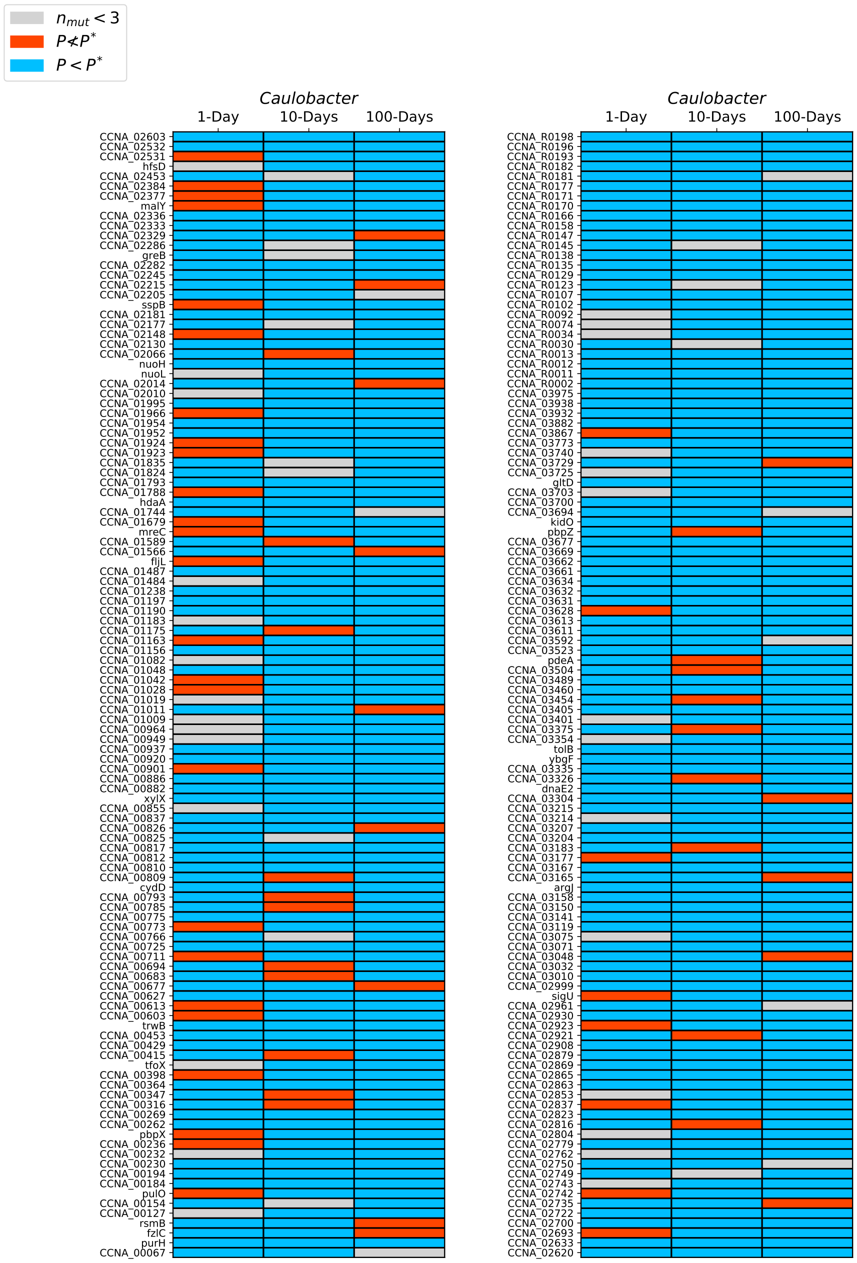


Figure S22: Visualization of all genes with an excess of non-synonymous mutations in more than one treatment for *Caulobacter*.

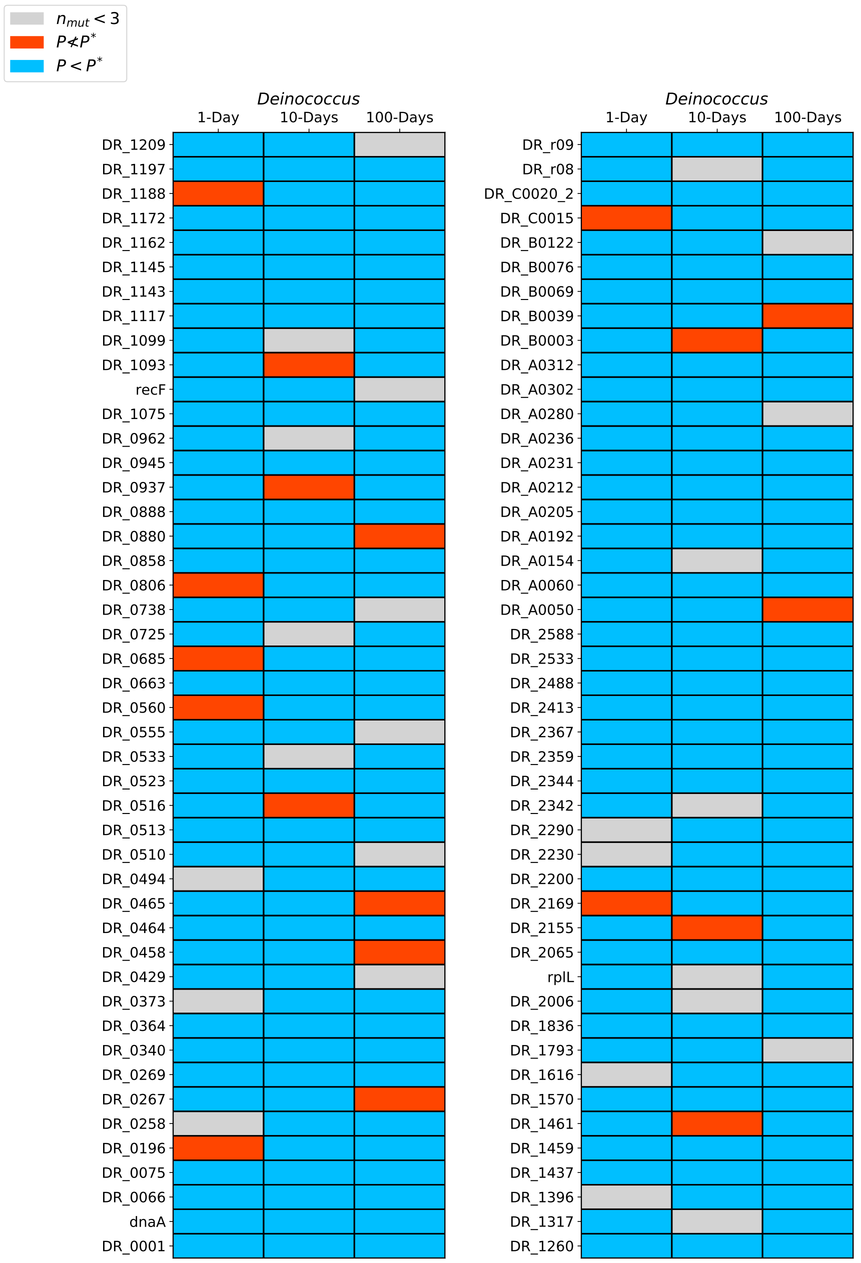


Figure S23: Visualization of all genes with an excess of non-synonymous mutations in more than one treatment for *Deinococcus*.

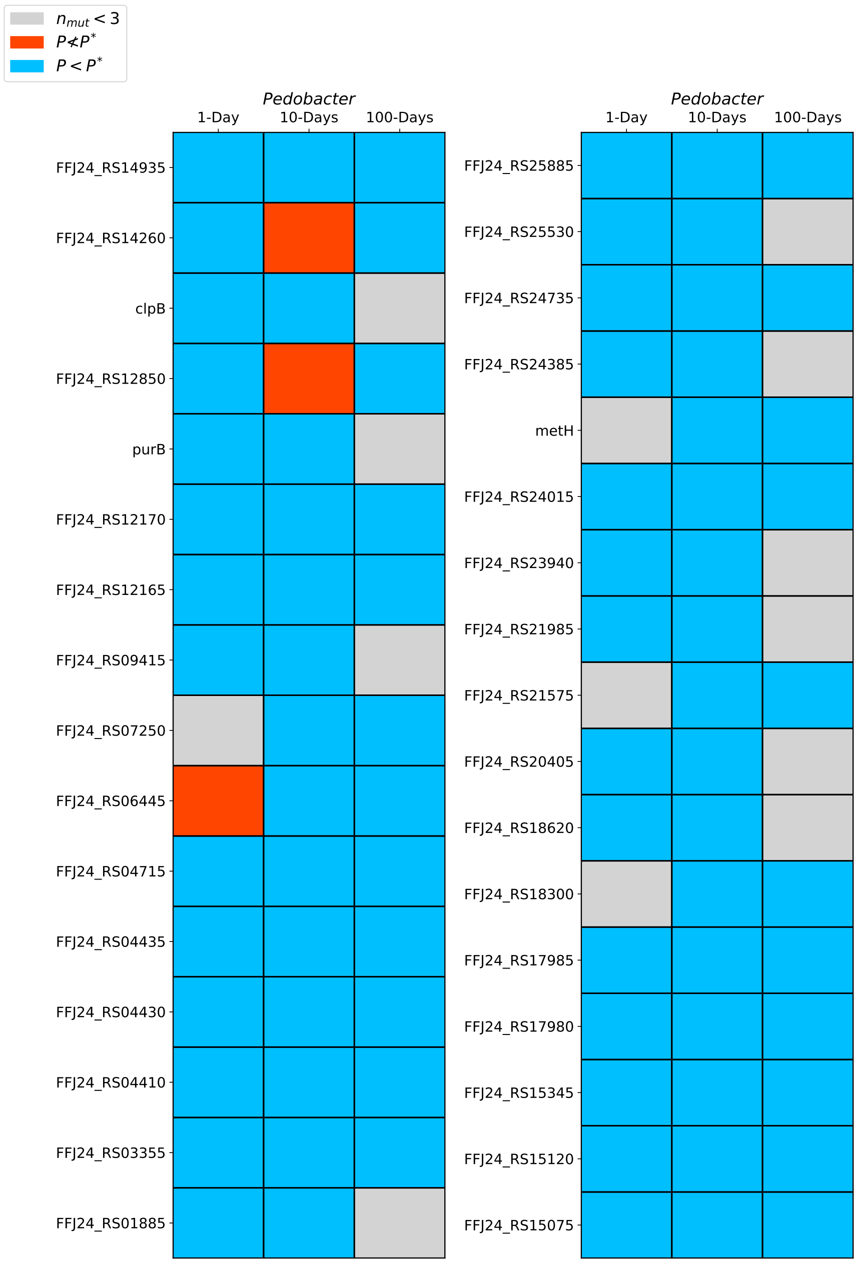


Figure S24: Visualization of all genes with an excess of non-synonymous mutations in more than one treatment for *Pedobacter*.

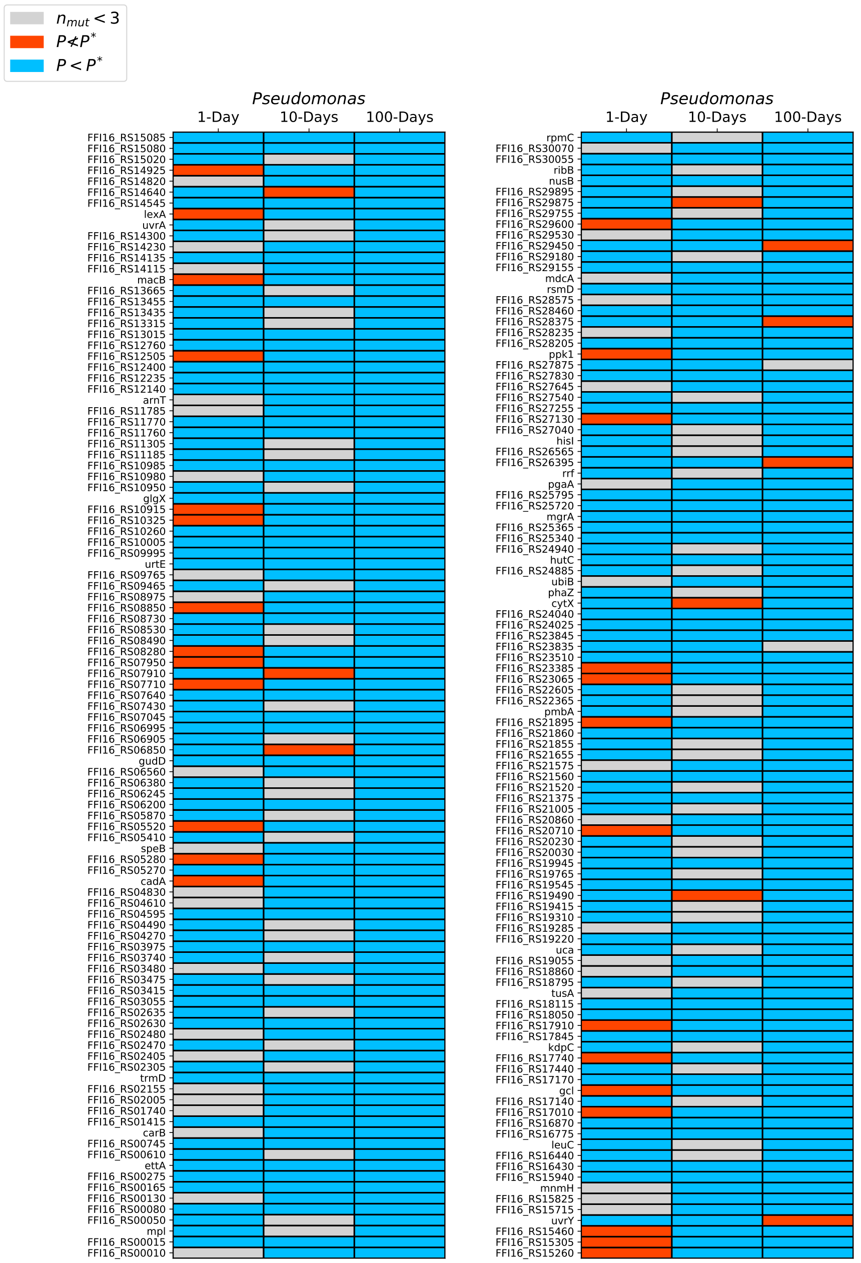


Figure S25: Visualization of all genes with an excess of non-synonymous mutations in more than one treatment for *Pseudomonas*.
